# Dynamic Regulation OF The Chromatin Environment By Ash1L Modulates Human Neuronal Structure And Function

**DOI:** 10.1101/2024.12.02.625500

**Authors:** Megha Jhanji, Joseph A. Ward, Calvin S. Leung, Colleen L. Krall, Foster D. Ritchie, Alexis Guevara, Kai Vestergaard, Brian Yoon, Krishna Amin, Stefano Berto, Judy Liu, Sofia B. Lizarraga

## Abstract

Precise regulation of the chromatin environment through post-translational histone modification modulates transcription and controls brain development. Not surprisingly, mutations in a large number of histone-modifying enzymes underlie complex brain disorders. In particular, the histone methyltransferase ASH1L modifies histone marks linked to transcriptional activation and has been implicated in multiple neuropsychiatric disorders. However, the mechanisms underlying the pathobiology of ASH1L-asociated disease remain underexplored. We generated human isogenic stem cells with a mutation in ASH1L’s catalytic domain. We find that ASH1L dysfunction results in reduced neurite outgrowth, which correlates with alterations in the chromatin profile of activating and repressive histone marks, as well as the dysregulation of gene programs important for neuronal structure and function implicated in neuropsychiatric disease. We also identified a novel regulatory node implicating both the SP and <u>Krüppel</u>-like families of transcription factors and ASH1L relevant to human neuronal development. Finally, we rescue cellular defects linked to ASH1L dysfunction by leveraging two independent epigenetic mechanisms that promote transcriptional activation. In summary, we identified an ASH1L-driven epigenetic and transcriptional axis essential for human brain development and complex brain disorders that provide insights into future therapeutic strategies for ASH1L-related disorders.

## INTRODUCTION

Post-translational modifications of histones, such as methylation, provide a mechanism to modulate gene expression broadly ^1–4^. The precise regulation of gene expression is essential for proper brain development. In fact, a large number of neuropsychiatric and neurodevelopmental disorders arise from mutations in enzymes that modulate the chromatin environment ^5^. The chromatin regulator *Absent, small, homeotic-like* (*ASH1L)* is a major genetic risk factor for autism spectrum disorders (ASD), Tourette syndrome (TS), and schizophrenia (SCZ) as determined by large-scale sequencing studies ^6–11^. To date, 136 disease variants in ASH1L have been associated with a wide spectrum of variable phenotypes in disorders of neuronal connectivity including ASD, attention-deficit/hyperactivity disorder (ADHD), TS, SCZ, intellectual disability (ID), and epilepsy ^6–16^. Knockout of ASH1L in mice results in a disorganized cortical architecture ^17^, growth retardation ^18^, synaptic defects, and neuronal hyperactivity ^19^. Taken together the human clinical studies and the animal studies suggest that ASH1L may play a key role in the establishment and maintenance of neuronal connectivity.

*ASH1L* encodes a histone methyltransferase of the Trithorax protein family that di-methylates lysine 36 on histone H3 (H3K36me2) ^20–22^ and has been associated with the tri-methylation of lysine 4 on histone H3 (H3K4me3). Both H3K36me2 and H3K4me3 are histone marks associated with transcriptional activation ^23^. In addition, H3K36me2 is proposed to sterically hinder the histone methyltransferase activity of Polycomb repressor complex 2 (PRC2) preventing it from tri-methylating lysine 27 on histone H3 (H3K27me3) ^20,24^, a mark associated with transcriptional repression ^4,25^. Therefore, loss of ASH1L could lead to unopposed widespread transcriptional repression by PRC2 on ASH1L target genes. Further, through its interaction with the CREB binding protein (CBP), the fly homolog of ASH1L might promote acetylation of H3K27 (H3K27ac), which could in turn oppose the catalytic activity of PRC2 ^26^. Additionally, inhibition of histone deacetylase activity (HDAC) by Vorinostat, an FDA approved cancer drug ^27^, rescued autism-like behaviors in a mouse mutant for ASH1L ^28^. Hence, ASH1L could modulate gene expression by directly and indirectly antagonizing the activity of PRC2 ^20,21,29^. However, whether the ASH1L/PRC2 axis regulates essential programs to build human neuronal connectivity remains poorly understood.

To model cellular and molecular phenotypes relevant to ASH1L-related disorders, we generated isogenic induced pluripotent stem cells (iPSCs) in which we engineered a pathogenic variant in ASH1L’s catalytic domain that has been associated with ASD, intellectual disability (ID), and epilepsy. Our results show that ASH1L dysfunction leads to impaired neuronal development, characterized by reduced neurite outgrowth and simplified neuronal arbors. Transcriptomic and epigenetic analysis of ASH1L mutant neurons suggest that ASH1L modulates gene regulatory programs essential for establishing and maintaining neuronal connectivity. This ASH1L-driven transcriptional and epigenetic node implicates gene programs involved in axonal development, neuronal migration, axon guidance, synaptic function, transcriptional regulation, and WNT signaling. Further, our analysis reveals strong links between ASH1L dysfunction and neuropsychiatric disorders such as ASD, epilepsy, and addiction while also suggesting potential connections to neurodegenerative disorders. Finally, we tested two rescue strategies that harness the power of modulating the chromatin environment using two FDA-approved small molecules that inhibit either PRC2 catalytic activity or HDAC activity. Albeit through different mechanisms, both strategies prevent transcriptional repression, promote transcriptional activation, and rescue the neuronal morphogenesis defects associated with ASH1L pathogenic variants. In summary, our studies suggest that an ASH1L/PRC2 axis and an ASH1L/histone acetylation axis regulates the development of human neuronal connectivity, and its disruption constitutes a novel mechanism underlying ASD and epilepsy.

## RESULTS

### Mutations in *ASH1L’s* catalytic domain lead to defects in neuronal arborization

In order to interrogate the function of ASH1L, we selected a pathogenic variant in ASH1L that targets its catalytic domain and is associated with ASD, severe intellectual disability (ID) and seizures ^7,30,31^ (**Fig.1A**). ASH1L-related pathology results from loss of function mutations causing a haploinsufficiency syndrome ^7,32^. To model ASH1L haploinsufficiency we used a previously published human iPSC line (11A) obtained from a healthy neurotypical male donor^33,34^ to introduce a heterozygous nonsense mutation, p.E2148* ^7,32^ (catalytic domain), in ASH1L using the CRISPR/CAS9 Ribonucleoprotein (RNP) system ^35^ ^36^ (**Fig.1A**, **Supplementary Fig. S1A and Supplementary Table S1)**. iPSCs containing the pathogenic mutation E2148* appear to maintain their pluripotency in culture, showing minimal spontaneous differentiation ^37,38^ as well as no significant changes in gene expression of pluripotency genes *OCT4, SOX2 and NANOG* (**Supplementary Fig. S1B-C and Supplementary Table S2**). Similarly, biochemical analysis of OCT4 and NANOG did not show significant changes in E2148* mutant iPSCs compared to control iPSCs (**Supplementary Fig. S1E**). However, we found decreased protein levels of SOX2 in E2148* mutant iPSCs (**Supplementary Fig. S1E**), which may suggest a novel role for ASH1L in the regulation of cell plasticity early in embryonic development ^39^.

**Figure 1.**
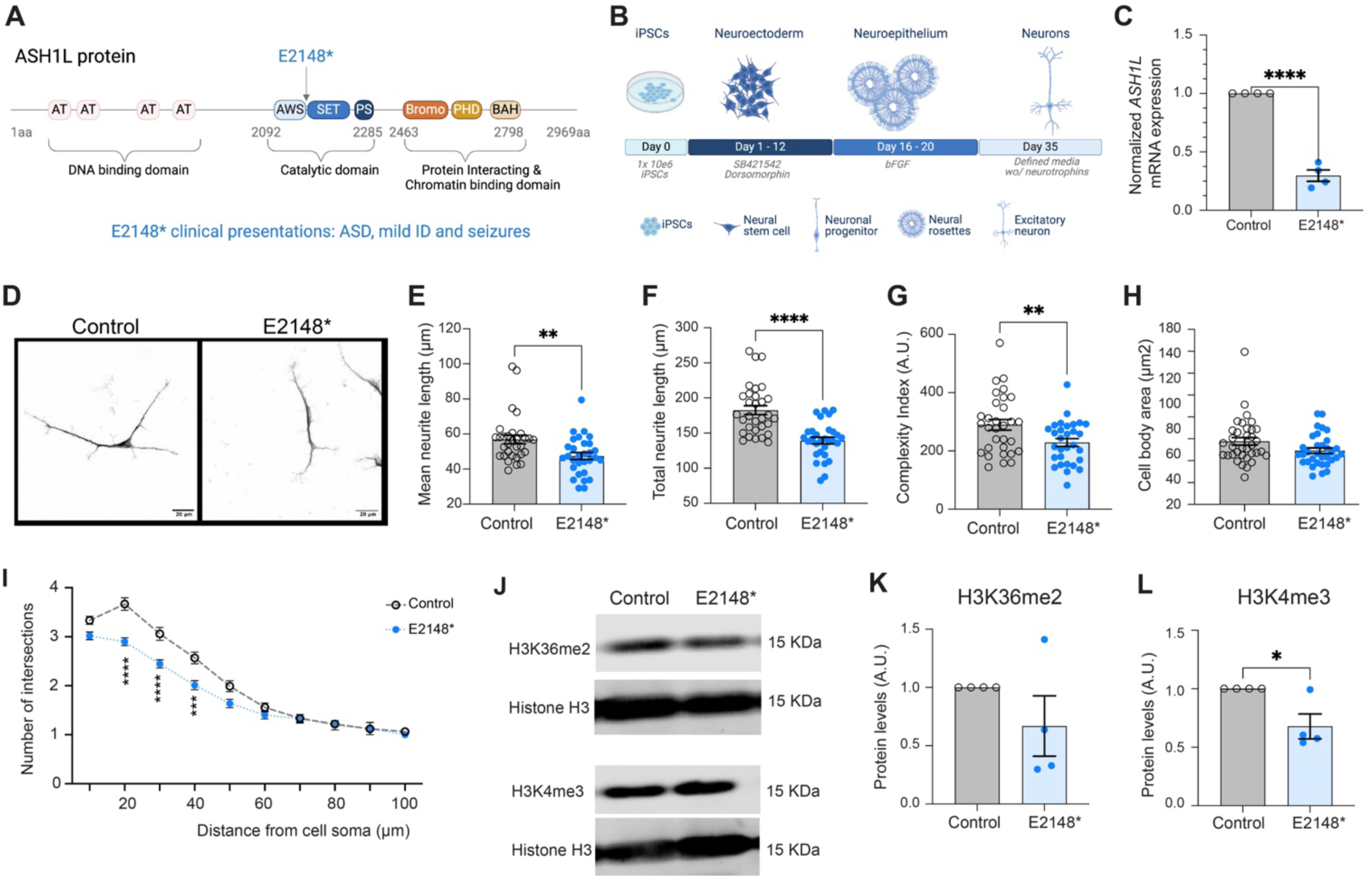
Neurite outgrowth and morphogenesis is altered in human neurons with pathogenic variant in the catalytic domain of ASH1L. **(A)** Diagram of ASH1L protein domains showing the location of the pathogenic variant E2148* (blue) in ASH1L catalytic domain and itsr associated clinical phenotypes. **(B)** Illustration depicts the dual SMAD inhibition protocol used to generate cortical excitatory human neurons. (**C**) ASH1L expression was quantified by qPCR using human neurons at day 35 of neuronal induction. Fold change is normalized to control. Bar represents the mean and individual measures from four independent experiments are shown for control (grey with open circles), and E2148* (light blue with solid blue circles). Samples were analyzed as a ratio of the control. Statistical analysis was conducted using unpaired t-test. **** P < 0.0001. (**D**) Representative images are shown for human neurons from control, and E2148* cultures at day 35 of neuronal induction. Neurons stained with MAP2 are shown in black and white for ease of viewing. Calibration bars represent 20µm. (**E-H**) Morphogenesis measures are shown for four independent experiments for control neurons (grey bar with open circles), and E2148* mutant neurons (light blue bars with solid dark blue circles). Individual points represent the average of 4 independent experiments, an average of 30 neurons were measured per experiment. (**E**) Mean neurite length is shown for control (n=124 neurons; 56.9 ± 2.41), and E2148* (n=118 neurons; 47.47 ± 1.99). Grouped statistical analysis was conducted using unpaired t-test, **P < 0.004. (**F**) Total neurite length is shown for control (n=124 neurons; 182.7 ± 6.39), and E2148* (n=118 neurons; 139.3 ± 4.66). Grouped statistical analysis was conducted using unpaired t-test, **** P < 0.0001. (**G**) Neuronal morphology analyzed by measuring the complexity index (see methods). Calculations were conducted after identifying outliers using the ROUT 1% method for control (n=115; 289.5 ± 18.21), and E2148* (n=112; 228.8 ± 13.42). Grouped statistical analysis was conducted unpaired t-test ** P < 0.0099. (**H**) Cell soma size was analyzed for three independent experiments by measuring the area for control (n=96; 77.67 ± 3.47), and E2148* (n=91; 69.15 ± 2.51). Statistical analysis was conducted using unpaired t-test P=0.056. (**I**) Sholl analysis was used to measure neuronal arborization. The number intersections away from the cell soma were measured every 10µm and are shown for control (open gray circles), and E2148* (solid dark blue circles) neurons from 10µm to 120µm. Statistical analysis was conducted using a mixed model effects *** P < 0.0006, and **** P < 0.0001. (**J-L**) Analysis of H3K36me2 and H3K4me3 levels on chromatin fraction for four independent experiments is shown for neurons at day 41 of neuronal induction. (**J**) Representative western blot shows H3K36me2, H3K4me3 and histone H3 for control, and E2148* neurons. H3 Histone marks were normalized to histone H3 levels for analysis. (**K**) H3K36me2 protein levels are shown for control (1± 0), and E2148* (0.67 ± 0.26). (**L**) H3K4me3 protein levels are shown for control (1± 0), and E2148* (0.68± 0.11). (K**-L**) Statistical analysis was conducted using unpaired t-test *P< 0.025. Not significant P value is not shown.

To investigate the neuronal phenotypes associated with ASH1L dysfunction, we differentiated iPSC lines into cortical excitatory neurons ^36,40^ using a dual SMAD inhibition approach ^36,40,41^ (**Fig. 1B**). Neurons were analyzed at day 35 of neuronal differentiation at which point the cultures are primarily composed of immature early born cortical excitatory neurons ^42^. Neurons with E2148* mutation showed reduced *ASH1L* mRNA expression by quantitative PCR (qPCR) compared to isogenic control neurons (**Fig.1C**). Based on our previous work in which human embryonic stem cell (ESC) derived neurons with knockdown of ASH1L showed reduced neuronal arborization ^40^, we analyzed the impact of mutating the catalytic domain of ASH1L on neuronal morphogenesis (**Fig. 1D-I**). We found that compared to control neurons, iPSC-derived cortical excitatory neurons ^36,41^ containing the E2148* pathogenic variant in *ASH1L* have reduced neurite outgrowth (**Fig. 1E-F**) and decreased arbor complexity by Sholl and complexity index analysis (**Fig.1G and 1I**), as we previously observed in ESC-neurons with ASH1L knockdown ^40^. However, the changes in morphology did not include a change in soma size as seen in ESC-derived neurons with knockdown for ASH1L ^40^ (**Fig. 1H**). Taken together, our data suggests that independent of genetic background, ASH1L plays a key role in the regulation of neuronal arborization during human brain development.

Differential histone methylation can have a profound impact on transcriptional activation or repression ^14^ which in turn could alter gene programs important for neuronal structure development and/or maintenance. ASH1L dimethylates histone H3 on lysine 36 (H3K36me2), which is found in transcriptionally active gene bodies ^21^. ASH1L has also been associated with the tri-methylation of lysine 4 on histone H3 (H3K4me3) ^43,44^, which is mainly enriched at transcription start sites (TSSs) and correlates with active gene expression. To investigate whether changes in H3K4me3 or H3K36me2 could be underlying the neuronal structure phenotypes, we measured total protein levels of H3K36me2 and H3K4me3 on neuronal chromatin fractions (**Fig.1J-L**), finding a significant decrease in H3K4me3 (**Fig.1J and 1L**) and a similar trend (albeit not significant) in H3K36me2 levels in the chromatin fraction (**Fig.1J-K**). In contrast, analysis of H3K36me2 and H3K4me3 in iPSC total cell lysates showed significant decrease of both H3K36me2 and H3K4me3 in E2148* mutant iPSC compared to controls (**Supplementary Fig. S1E**). Multiple histone methyltransferases besides ASH1L have been previously associated with the deposition of both H3K36me2 and H3K4me3 ^5^. Therefore, the bulk changes observed in iPSCs for both H3K36me2 and H3K4me3 and in neurons primarily in H3K4me3 suggest an important function for ASH1L during development.

### *ASH1L* modulates broad transcriptional programs important for neuronal structure and function by regulating gene expression and transcription efficiency

Based on the biochemical bulk analysis of ASH1L-related histone marks, we posited that pathogenic variants in ASH1L might lead to widespread alterations in the neuronal transcriptome. We performed strand-specific bulk RNA-seq ^45^ on human neurons differentiated for 35 days to correlate with morphological studies (**Supplementary Fig. S2A**). We obtained an average of 100 million reads per sample that were mapped to the human genome and were analyzed for differential gene expression^45^ (**Fig. 2**, **Supplementary Fig. S2, and Supplementary Table S3**).

**Figure 2.**
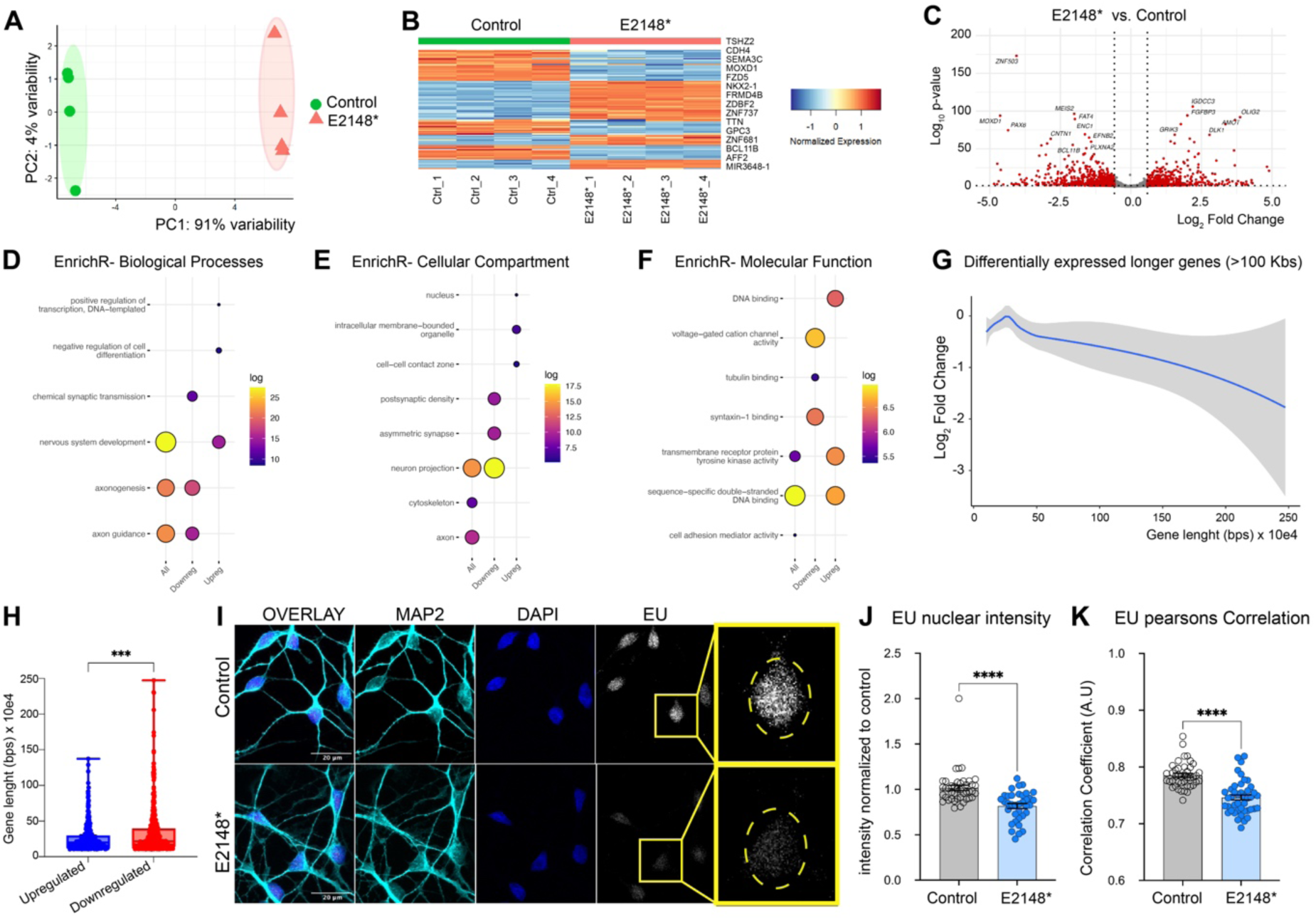
ASH1L modulates transcriptional programs associated with neuronal structure and function, as well as transcription of longer genes and transcription efficiency. (**A**) PCA plots shows biological replicates (n=4) for control (green), and E2148* (salmon) neurons RNA seq experiments. (**B**) Heatmap shows top 100 DEGs for control (green), and E2148* (salmon) neurons at day 35 (n=4 biological replicates). The top 15 DEGs are listed. (**C**) Volcano plots showing DEGs in the heterozygous E2148* mutant iPSC-derived neurons. Log_2_ fold changes (LFC) gene expression (x-axis) and -log_10_ adjusted P values (y-axis) generated from DESeq2 differential gene expression analysis are shown. Vertical dotted lines represent 0.58 LFC (1.5 FC) and horizontal dotted line shows adjusted P=0.05. Significant DEGs are shown in red with the top 20 labelled in the plot. (**D-F**) Functional enrichment analysis by EnrichR for biological process (**D**), cellular compartment (**E**), and molecular function (**F**) show enrichment for all DEGs, upregulated and downregulated DEGs in E2148* mutant neurons vs. control neurons. Circle size represents the number of DEGs in that category and the color represents the adjusted P value. (**G**) Correlation of gene length to fold change analyzed for all significant DEGs in E2148* (blue line) mutant neurons. Grey shade shows the variability across samples. (**H**) Analysis of gene length in upregulated (blue) and downregulated (red) DEGs for E2148* neurons. (**I**) Analysis of de novo transcription by EU click chemistry at day 41 of neuronal differentiation. Representative images of human neurons that incorporated EU (gray), stained with neuronal marker MAP2 (cyan) and nuclear marker DAPI (blue) are shown for control (top row), and E2148* (bottom row). Enlarged nuclei stained with EU is shown. Calibration bars are 20µm. (**J-K**) Measurements of EU incorporation are shown for control neurons (grey bars with open circles), and E2148* (light blue bars with solid deep blue circles) mutant neurons. Mean and standard error are shown with individual dots representing the average of individual measures for five independent experiments. (**I**) Pearsons’ correlation coefficient analysis is shown for five independent experiments for control (n=187; 0.785 ± 0.003), and E2148* (n=115; 0.746 ± 0.005) neurons. (**J**) EU nuclear intensity normalized to control is shown for five independent experiments for control (n=187; 1.017 ± 0.029), and E2148* (n=115; 0.817 ± 0.027) neurons. (**I-J**) Grouped data analyzed using unpaired t test with Welch’s correction, ****P < 0.0001. Not significant P values are not shown.

Our analysis of coding and non-coding transcripts showed that the ASH1L mutant neurons formed a distinct cluster, separate from controls, suggesting widespread transcriptional changes (**Fig 2A**). Differential expression analysis identified 2475 differentially expressed genes (DEGs) (**Fig. 2B-C**, **and Supplementary Table S3**). Functional enrichment analysis by *EnrichR* showed a significant overrepresentation of biological processes associated with axon guidance, axonogenesis, nervous system development, transcriptional regulation and synaptic function (**Fig. 2D**, **and Supplementary Table S4**). Cellular compartment analysis showed enrichment for genes encoding proteins localized to the neuronal projections, axons, synaptic structures, cell-cell contact zones, and the nucleus (**Fig. 2E and Supplementary Table S4**). Molecular function analysis further supported a role for ASH1L in the modulation of gene programs important for synaptic and neuronal transmission pathways (voltage gated cation channel; syntaxin-1 binding activity), and regulation of transcriptional programs (DNA binding) (**Fig. 2F and Supplementary Table S4**). Additional gene ontology (GO) analysis of biological processes confirmed the importance of ASH1L in the regulation of neuronal structure and synaptic function (**Supplementary Fig. S2B-C and Supplementary Table S4**), and revealed the upregulation of genes associated with the canonical WNT signaling pathway in human neurons (**Supplementary Fig. S2D and Supplementary Table S4**). GO analysis of molecular function revealed that downregulated DEGs clustered within synaptic function pathways (i.e. voltage gated ion channel activity, glutamate receptor binding) and translation-related processes (i.e. structural constituent of ribosome). In contrast, upregulated DEGs highlight roles in cell signaling (i.e. extracellular matrix, transmembrane receptor protein kinase activity), cell adhesion and chromatin dynamics (**Supplementary Fig. S2E-F and Supplementary Table S4**). Overall, our results indicate a disruption of pathways important for neuronal development and synaptic function, emphasizing the impact of mutations in the catalytic domain of ASH1L on neurodevelopmental processes.

Given the enrichment of synaptic and axonal genes among the DEGs modulated by ASH1L and the known dysregulation of transcription of long genes (>100 kbs) in neurodevelopmental disorders ^46,47^, we analyzed the correlation between DEGs and gene length using Loess curve regression. E2148* mutant neurons showed a length-dependent decrease in gene expression, with the longer genes showing a more significant reduction in expression (**Fig. 2G and Supplementary Table S3**), which suggests a potential impairment in transcription efficiency, that could contribute to the reduced expression of synaptic and axonal genes in the ASH1L mutant neurons. Additionally, comparing gene length between downregulated and upregulated DEGs in E2148* mutant neurons revealed a preferential downregulation of longer genes (**Fig. 2H**). To further assess changes in transcription efficiency, we performed a 5-ethynyl uridine (EU) incorporation assay in ASH1L mutant neurons. We measured nascent RNA synthesis in day 39 neurons by incorporating the uridine analog EU at 100 μM for 3 hours into newly transcribed RNA, followed by detection using click chemistry **(Fig. 2I-K)**. We found that E2148* mutant neurons showed reduced EU incorporation by nuclear intensity and Pearson’s correlation coefficient analysis **(Fig. 2J-K)**, indicating decreased transcription efficiency, which could be due to increased gene repression in the absence of ASH1L catalytic activity. Finally, based on previously published chromatin immunoprecipitation sequencing data (ChIP-seq) ^48^ we analyzed transcription factor binding targets across upregulated and downregulated DEGs in E2148* mutant neurons. We found a transcriptional regulatory node essential for neuronal development that included major transcription factors such as DEAF1, NEUROD2, and REST (**Supplementary Table S5**).

### Disruption of *ASH1L* in human neurons leads to distinct changes in alternative splicing

We previously showed that knockdown of ASH1L in ESC-derived human neurons alters three key splicing isoforms of the BDNF receptor TrkB (encoded by the *NTRK2* gene) that are important for neuronal arborization ^40^. ASH1L-mediated H3K36me2 acts as a substrate for trimethylation at lysine 36 on histone H3 (H3K36me3) ^49^, and ASH1L interacts with MRG15 which acts as a hub to recruit splicing proteins to H3K36me3 ^50^ sites in the chromatin. To determine if changes in isoform usage were a generalized effect associated with deficits in ASH1L’s catalytic activity, we conducted a differential analysis of transcript usage (DTU). We found significant alterations in DTU across 57 genes, including those important for neurodevelopment (**Fig. 3**, **Supplementary Figure S3, and Supplementary Table S6**). Although we did not find specific splicing events being altered (**Fig. 3B**) or a large convergence of genes with DTU in specific pathways, we found that the most significant DTU events in ASH1L mutant neurons resulted in production of distinct isoforms missing key functional domains or with different sensitivities to nonsense-mediated decay (NMD) (**Fig. 3C-E**, **Supplementary Fig. S3A-B and Supplementary Table S6**). For example, E2148* mutant neurons showed increased usage of an isoform of the chromatin remodeler SMARCA4 that lacks ATPase activity **(Fig. 3C-E**). Although the overall gene expression of *SMARCA4* (**Fig. 3D**) and its isoform expression (**Fig. 3E**) did not show significant changes, we observed notable differences in *SMARCA4* transcript usage (**Fig. 3C**). Similarly, we detected a significant DTU in *AFF2* (**Supplementary Fig. S3A**), which encodes a protein involved in mRNA splicing and processing ^51^. Specifically, we observed decreased usage of an isoform of *AFF2* missing several key AF4 domains important for protein-protein interactions and its role in transcriptional elongation. While significant changes in DTU for both *AFF2* and *SMARCA4* is in one of their protein-coding isoforms, *TARDBP*, a regulator of transcription and RNA stability ^52^ shows decreased DTU in a non-coding isoform that is insensitive to NMD (**Supplementary Fig. S3B**). AFF2 and SMARCA4 have both been associated with neurodevelopmental disorders that present with ASD ^53^ ^53^ while *TARDBP* encodes TDP-43 which is known for its association with amyotrophic lateral sclerosis (ALS) ^52^. However, while neither *SMARCA4* nor *TARDBP* showed significant changes in gene expression, *AFF2* is among the significantly upregulated DEGs in ASH1L mutant neurons. Taken together our previous work and our current findings suggest that ASH1L may regulate the transcriptome at multiple levels, either by differentially regulating gene expression and splicing or by influencing both in a subset of genes, such as *NTRK2* and *AFF2*.

**Figure 3.**
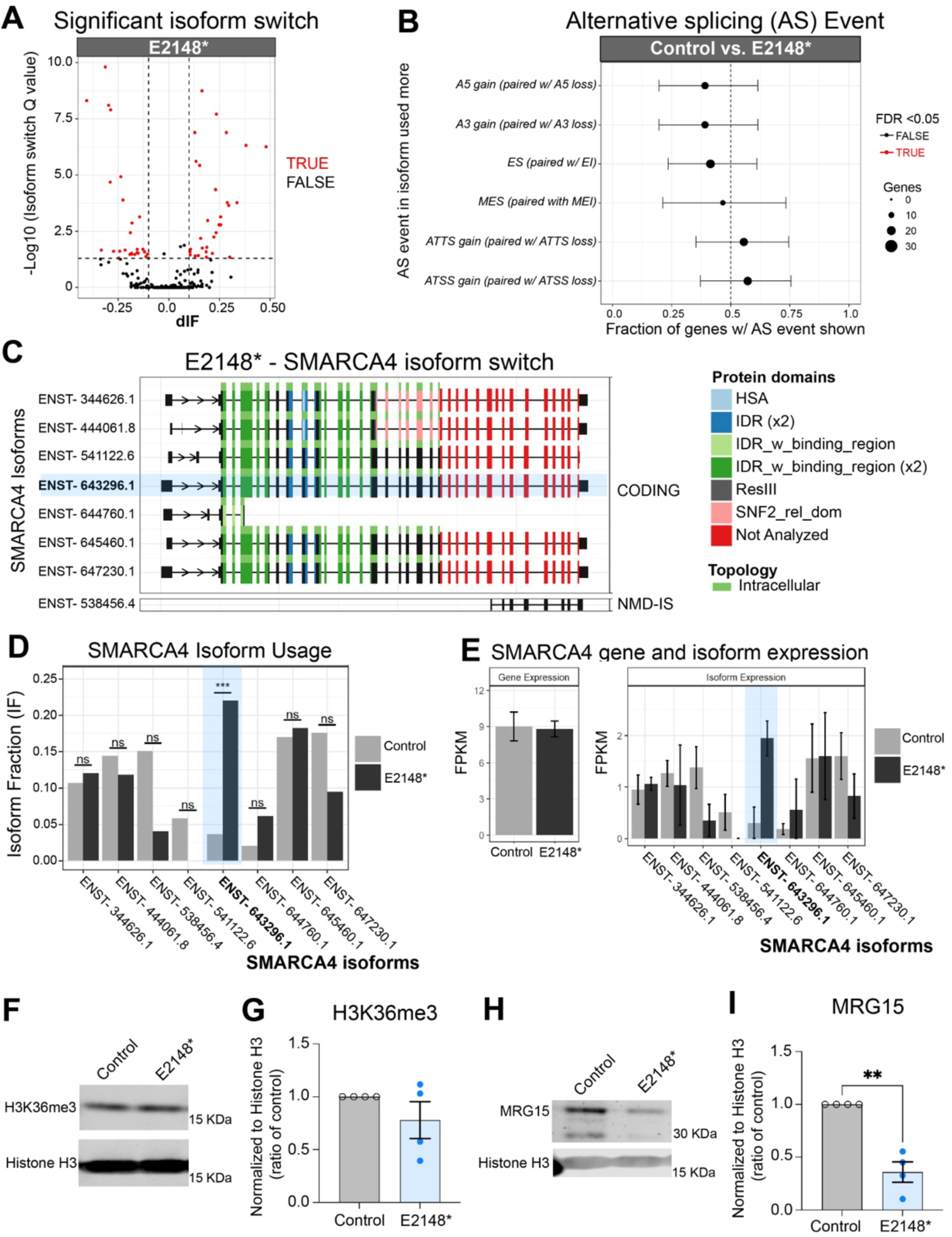
ASH1L regulates differential transcript usage in human neurons. (**A**) Significant isoform switching events are shown in red raising above adjusted p value (Q) plotted as the -Log_10_ (Q) (y axis) and the differential isoforms (dIF) (x axis) for E2148* mutant neurons compared to control neurons. (**B**) Types of alternative splicing events were analyzed for mutant neurons with respect to control neurons by plotting the AS event present in the most used isoform (y axis) vs. the fraction of genes with AS event per isoform (x axis). (**C**) Graph shows all the coding and non-coding isoforms for SMARCA4. (**D**) Analysis of isoform usage for SMARCA4 in control (grey) and E2148* mutant neurons (black) compared to control neurons (grey), with the most significant changed isoform highlighted in blue. (**E**) Analysis of SMARCA4 gene and isoform expression in control (grey) and E2148* mutant neurons (black). (**F**) Representative image shows levels of H3K36me3 in chromatin fraction for control, and E2148* mutant neurons. (**G**) Analysis of H3K36me3 across four independent experiments for control (grey bar open circles; 1 ± 0), and E2148* (light blue bar and dark blue solid circles; 0.78 ± 0.17) mutant neurons. (**H**) Representative image shows levels of MRG15 in chromatin fraction for control, and E2148* mutant neurons. (**I**) Analysis of MRG15 across four independent experiments for control (grey bar open circles; 1 ± 0), and E2148* (light blue bar and dark blue solid circles; 0.36 ± 0.097) mutant neurons. Statistical analysis (**G and I**) conducted with unpaired t-test with Welch’s correction. ** P< 0.008, not significant P values are not shown.

To explore the mechanisms underlying DTU in ASH1L mutant neurons, we examined the levels of tri-methylated H3K36 (H3K36me3) and MRG15. H3K36me3 can act as a hub to recruit splicing factors^54^, and while ASH1L does not directly deposit H3K36me3, it provides the H3K36me2 substrate necessary for its production ^55^. MRG15 binding to H3K36me3 mediates the chromatin interaction with splicing factors and ASH1L is an interactor of MRG15 ^50^. We detected no significant changes in bulk H3K36me3 levels (**Fig. 3F-3G**) but observed a significant reduction in MRG15 protein levels (**Fig. 3H-3I**). Taken together, our findings suggest that DTU observed in ASH1L mutant neurons might be partly driven by reduced MRG15 activity rather than alterations in H3K36me3 levels.

### *ASH1L* controls transcriptional programs associated with multiple neurodevelopmental and neuropsychiatric disorders

Our functional enrichment studies underscored the importance of ASH1L in the development of neuronal circuitry. Defects in neuronal connectivity can lead to an array of neuropsychiatric and/or neurodevelopmental disorders. We sought to define whether alterations in the human neuronal transcriptome associated with ASH1L dysfunction could relate to the pathogenesis of brain disorders by examining disease association across non-coding and coding transcripts. Based on the preferential downregulation of longer genes in ASH1L mutant neurons and the correlation of ASD-risk with longer gene lengths^46^, we compared our dataset with a curated database of ASD-risk genes. Comparison with the Simons Foundation Autism Research Initiative (SFARI) database for ASD-risk genes ^15^ showed that 27% of the known ASD-risk genes are regulated by ASH1L in human neurons (**Fig. 4A**). Further, analysis of all DEGs using *DisGeNET* showed a strong association with cortical dysplasia, global developmental delay, seizures, autistic behaviors and Rett syndrome (**Fig. 4B and Supplementary Table S7**). Gene network analysis of DEGs present in the autistic behavior category showed an enrichment for genes that have high-risk variants for ASD, including genes like ARID1B, ASXL3, and AUTS2 (**Fig. 4C**). Further, the autistic behavior gene network contained several chromatin regulators (KDM5B, KMT5B and MEIS2) as well as transcriptional regulators (FOXP2, TCF4 and ZNF462) (**Supplementary Fig. S4A-B**). Interestingly, examination of downregulated DEGs by *DisGeNET* showed recurrent association with different forms of epilepsy in addition to cortical dysplasia and global developmental delay (**Fig. 4D and Supplementary Table S7**). Epilepsy and ASD are often co-morbid and this is shown by analysis of epilepsy gene networks, which contained several genes that have high-risk variants associated with ASD, such as KCNQ2, CNTNAP2, and GRIN2B (**Fig. 4E**). Additional analysis using OMIM disease and KEGG pathways highlight the known role of ASH1L in cancer pathogenesis, but also reveal a novel association with neurodegenerative disorders, addiction and circadian entrainment (**Supplementary Fig. S4C-D and Supplementary Table S7**).

**Figure 4.**
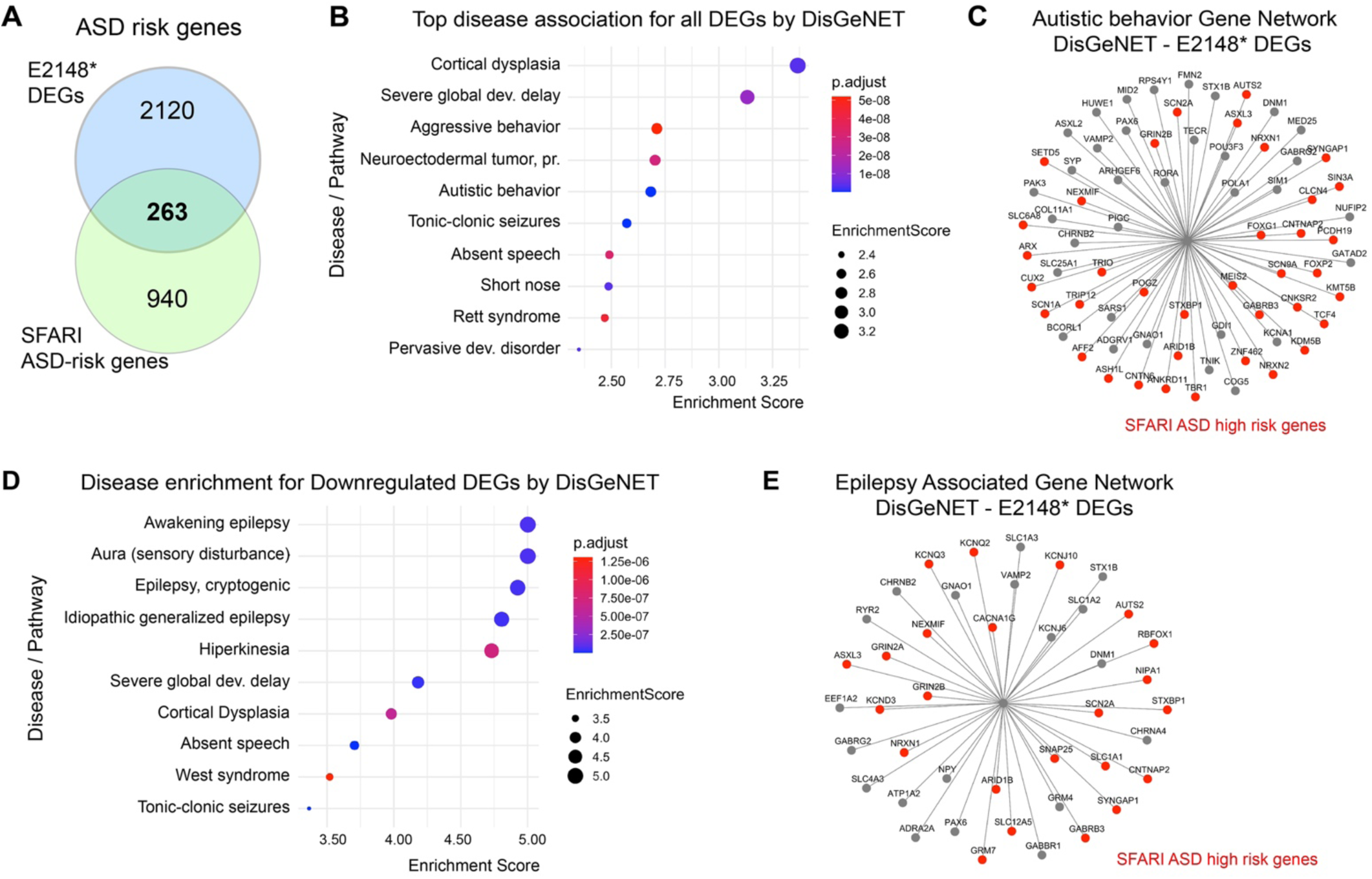
ASH1L modulates a disease nodule associated with autism and epilepsy. (**A**) Comparison of our DEG dataset with SFARI ASD gene database identified 263 DEGs that are ASD risk genes. Venn diagram shows overlapping ASD risk genes between the E2148* mutant neurons (blue) with SFARI database (green). (**B**) Disease association analysis of all DEGs using DisGeNET. The solid black circles indicate the enrichment score for each disease, while the color gradient represents the adjusted P values, with red indicating lowest value, and blue indicating highest value. (**C**) Starburst graph display the DEGs associated with autistic behavior shown in panel B. ASD-risk genes with high risk variants are shown in red. (**D**) Disease association analysis of only the downregulated DEGs using DisGeNET. The solid black circles indicate the enrichment score for each disease, while the color gradient represents the adjusted P values, with red indicating lowest value, and blue indicating highest value. (**E**) Starburst graph display the DEGs associated with epilepsy categories shown in panel D. ASD-risk genes with high risk variants are shown in red.

### Neuronal *ASH1L* modulates H3K36me2 and H3K4me3 occupancy in a genome-wide manner

The widespread changes in gene expression observed in human neurons with ASH1L dysfunction could be driven by alterations in post-translational histone modifications. H3K36me2 is a hallmark of ASH1L catalytic activity, and in non-neuronal cells, ASH1L has been associated with H3K4me3 methylation ^43,44^. We investigated genome-wide changes in H3K36me2 and H3K4me3 methylation status using Cleavage Under Targets and Tagmentation sequencing (CUT&Tag) ^56^ with validated CUT&Tag antibodies ^22,57,58^ ^59^ (**Fig.5**, **Supplementary Fig. S5-S6 and Supplementary Table S8-S9**). Heatmaps for the CUT&Tag peaks across the genome show a genome-wide decrease in H3K36me2 levels in mutant neurons compared to controls, whereas the H3K4me3 levels showed only a subtle reduction (**Fig.5A-B and Supplementary Table S8-S9**). While examining the distribution of differential peaks across gene regions (**Fig. 5C-E and Supplementary Fig. S5**), we found that H3K36me2 differential peaks (1034 peaks) were spread across gene bodies primarily in introns (35.4%) and exons (16.3%) in E2148* mutant neurons (**Fig. 5C and 5E**, **and Supplementary Fig. S5**). In contrast, the H3K4me3 differential peaks (3149 peaks) were primarily concentrated near TSSs with 20.5% at promoter regions in the E2148* mutant neurons (**Fig. 5D-E and Supplementary Fig. S5**).

**Figure 5.**
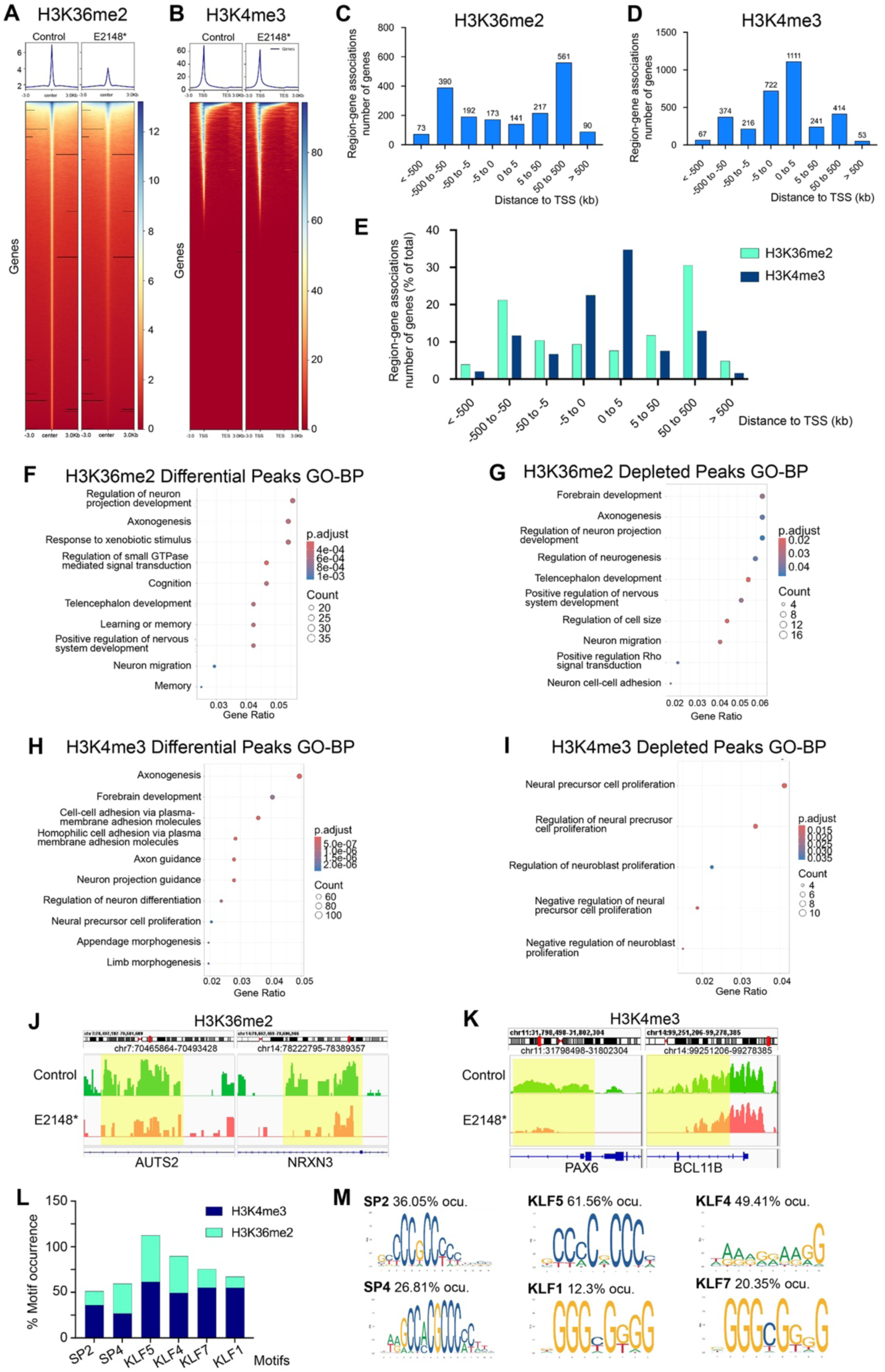
H3K4me3 and H3K36me2 profiling in human neurons with pathogenic variants in ASH1L. (**A-B**) Heatmaps showing all differential peaks aligned to the center within a 6kb region for H3K26me2 peaks (**A**) and aligned to transcription start site (TSS) for H3K4me3 peaks (**B**) for E2148* neurons is shown as an average of four independent experiments. (**C-E**) Histograms show the distribution of differential peaks gene-region associations in E2148* mutant neurons plotted as a function of the distance from the Transcription start site (TSS) vs. the number of genes per region for H3K36me2 (**C**), H3K4me3 (**D**) differential peaks. (**E**) The percentage of genes per region are shown for both H3K36me2 (light green) and H3K4me3 (dark blue). (**F-G**) Gene enrichment analysis by Gene Ontology (GO) for biological process (BP) for H3K36me2 differential peaks (**F**) and depleted peaks (**G**) in E2148* mutant neurons. (**H-I**) Gene enrichment analysis by Gene Ontology (GO) for biological process (BP) for H3K4me3 differential peaks (**H**) and depleted peaks (**I**) in E2148* mutant neurons. (**F-I**) The number of genes per category is shown by the size of the circles and the statistical significance is shown as the adjusted P value by the color hue. (**J-K**) Visualization of chromatin profiling for specific gene targets comparing control (green) vs. E2148* (salmon) mutant neurons for H3K26me2 (**J**) and H3K4me3 (**K**). Most differential peaks are highlighted in yellow. (**L**) Motif analysis of differential peaks is shown as motif occurrence percentage that are overlapping between H3K36me2 differential peaks (light green) and H3K4me3 differential peaks (dark blue) in E2148* mutant neurons. (**M**) Motif consensus sequence is shown for all overlapping transcription factors identified by motif analysis across H3K36me2 and H3K4me3 differential peaks.

Functional enrichment analysis of H3K36me2 and H3K4me3 differential peaks associated with gene encoding regions (**Fig. 5F-I**, **Supplementary Fig. S6, and Supplementary Table S10**) showed convergence across brain region specification (i.e. telencephalon and forebrain development) and neuronal connectivity (i.e. axonogenesis, neuron projection morphogenesis, asymmetric synapse, and postsynaptic density) (**Fig. 5F**, **5H, Supplementary Fig. S6, and Supplementary Table S10**). Interestingly, molecular function analysis showed that H3K4me3 differential peaks were enriched in processes related to transcriptional regulation, while H3K36me2 differential peaks showed a strong association with synaptic function (i.e. ion channel activity, calmodulin binding) (**Supplementary Fig. S6 and Supplementary Table S10**). To further interrogate the extent to which impairing ASH1L’s catalytic function alters the epigenome, we conducted functional enrichment analysis on enhanced and depleted differential peaks for H3K36me2 and H3K4me3 histone marks (**Fig. 5G**, **5I, Supplementary Fig. S6 and Supplementary Table S10**). Biological process analysis identified a distinct signature associated with the regulation of neural precursor proliferation across H3K4me3 depleted peaks (**Fig. 5I and Supplementary Table S10**); while H3K36me2 depleted peaks showed an association with the regulation of neurogenesis and neuronal migration (**Fig. 5G and Supplementary Table S10**). Molecular function analysis of H3K4me3 depleted peaks highlighted pathways important for synaptic function, such as cyclic AMP and voltage-gated ion channels (**Supplementary Fig. S6 and Supplementary Table S10**). In contrast, molecular function analysis of enhanced peaks highlighted an association with WNT signaling for H3K4me3 peaks and with small GTPase signal transduction for H3K36me2 peaks (**Supplementary Fig. S6 and Supplementary Table S10**).

Visualizing the differential peaks across specific target genes showed not only a reduction of peaks but also, in some cases, changes in peak distribution across gene bodies in mutant neurons compared to control neurons (**Fig. 5J-K**). These changes in histone methylation across gene bodies correlated with the gene expression findings; for instance, high-risk ASD genes such as AUTS2 and NRXN3 showed reduced expression and corresponding decreases in H3K36me2 across intronic regions in ASH1L mutant neurons (**Fig. 5J and supplementary Fig. S5H**). Similarly, significant decreases in H3K4me3 levels were observed in key neurodevelopmental transcription factors like PAX6 (neuronal progenitor marker), and BCL11B (gene encoding CTIP2 layer V transcription factor) with parallel reductions in mRNA levels in both genes by RNAseq (**Fig. 5K and Supplementary Fig. S5I**).

Finally, to identify potential regulomes affected by ASH1L, we conducted motif enrichment analysis across H3K36me2 and H3K4me3 differential peaks (**Fig. 5L-M and Supplementary Table S11**). Binding sites were preferentially enriched for SP family of transcription factors (SP2 and SP4) and Krüppel-like factors (KLF1, KLF4, KLF5 and KLF7) motifs across differential peaks for both histone marks (**Fig. 5L-M**) ^60,61^. The SP family of transcription factors is involved in the regulation of transcriptional elongation and has important functions during brain development ^62^. Specifically, SP2 is implicated in the proliferation of neuronal precursor cells ^63^, while SP4 contributes to the dendritic arbors patterning in the cerebellum ^64^. Similarly, the KLF family is integral to a regulatory network that modulates neuronal projection growth ^65,66^. These findings align with the functional enrichment signatures observed for H3K36me2 and H3K4me3 differential peaks, highlighting the involvement of both the SP and KLF regulatory nodes in neurogenesis, neuronal morphogenesis and structural development, critical developmental processes associated with ASH1L function.

### *ASH1L* dysfunction impacts PRC2-driven epigenetic mechanisms in human neurons

H3K27me3 is a hallmark of PRC2 activity, and is enriched at promoter regions of silenced genes ^25^. Both H3K36me2 and, in some instances, H3K4me3 marks are proposed to block the deposition of repressive H3K27me3 marks by PRC2 ^3,20^. We posit that ASH1L dysfunction could lead to unopposed PRC2 activity, increasing H3K27me3 levels and silencing essential neuronal genes. Using a CUT&Tag assay we identified broad genome-wide alterations of H3K27me3 occupancy in E2148* mutant neurons (**Fig. 6A**, **Supplementary Fig. S7 and Supplementary Table S12**). These changes were distributed across the gene bodies, with 3416 differential peaks in the *ASH1L* mutant neurons located 5-500 kb up or downstream from the TSS (**Fig. 6B and Supplementary Fig. S7A-B**), primarily within intronic regions (31.7%) (**Supplementary Fig. S7C**). We also find that between 1 to 3 genes per differential peak were contained across 3416 differential peaks (**Fig. 6C**). Visualization of H3K27me3 differential peaks showed an increase in peak size and distribution across genes associated with axon guidance, synaptic function, and neuronal differentiation (**Fig. 6D**). We examined PRC2 target genes, such as SEMA6D, CACNA2D3, STK33 and observed an increase in repressive PRC2-dependent methylation along with reduced mRNA expression, particularly for CACNA2D3 (**Fig. 6D and Supplementary Table S3)**. This inverse relationship between H3K27me3 levels and gene expression was also seen in genes like GRIN2B (NMDA receptor subunit) and SEMA5A (axon-guidance), both of which showed increased methylation and decreased expression in the ASH1L mutant neurons (**Fig. 6D and Supplementary Table S3)**.

**Figure 6.**
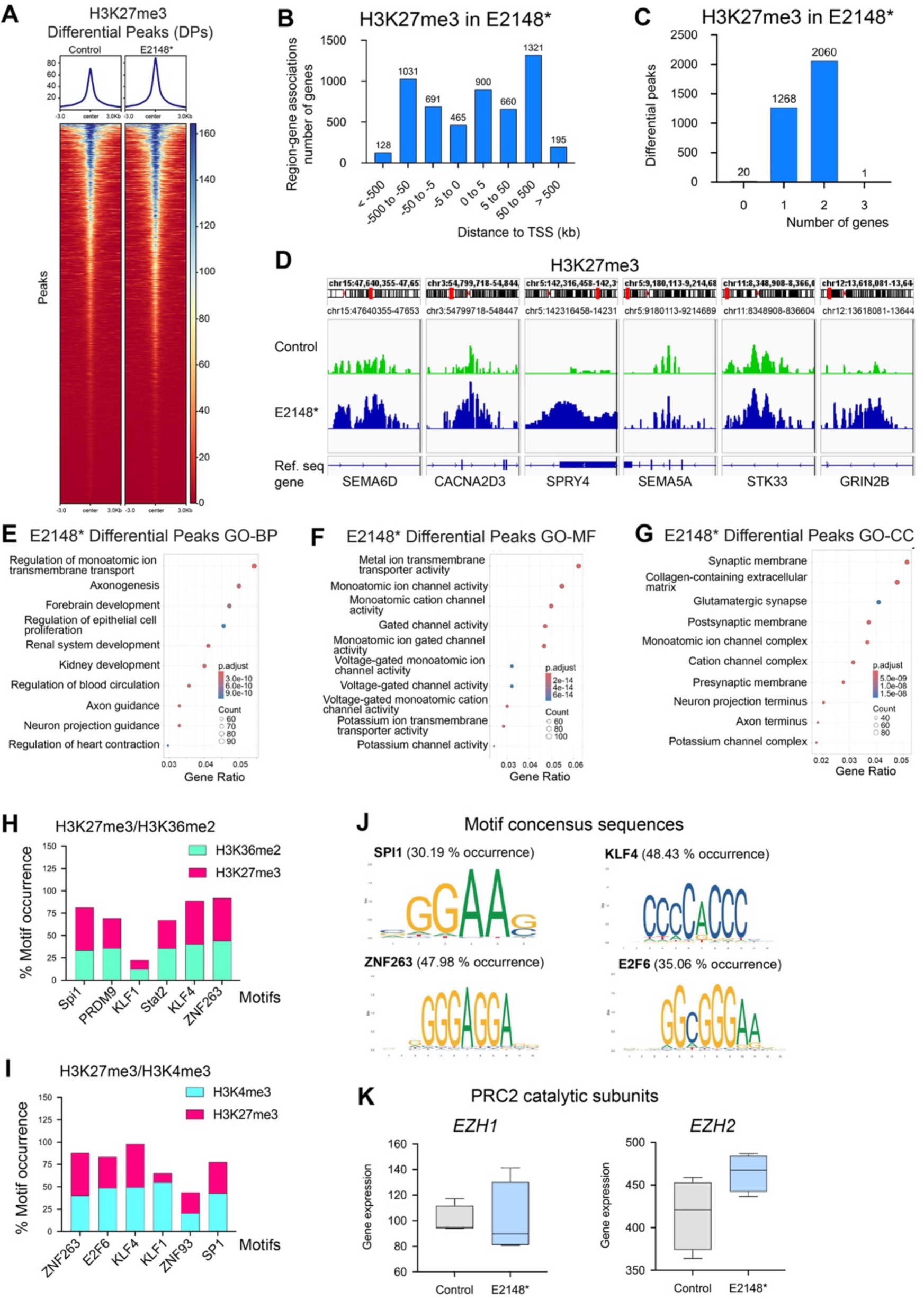
H3K27me3 chromatin profile is altered in human neurons with E2148* mutation in ASH1L. (**A**) Heatmaps showing all differential peaks aligned to the center within a 6kb region for H3K27me3 differential peaks for control, and E2148* neurons in four independent experiments. (**B**) Histograms show the distribution of H3K27me3 differential peaks gene-region associations as a function of the distance from the Transcription start site (TSS) in E2148* (blue bars) mutant neurons. (**C**) Histogram shows the number of genes per H3K27me3 differential peaks in E2148* (blue bars) mutant neurons. (**D**) Visualization of chromatin profiling for specific gene targets comparing control (green) vs. and E2148* (blue) mutant neurons for H3K27me3 differential peaks. (**E-G**) Functional enrichment analysis by Gene Ontology (GO) for Biological Process (BP) (**E**), Molecular function (MF) (**F**), and Cellular compartment (CC) (**G**), for H3K27me3 differential peaks in E2148* mutant neurons. The size of the genes per category is shown by the size of the circles and the statistical significance is shown as the adjusted p-value by the color hue. (**H-J**) Motif analysis of differential peaks in E2148* mutant neurons is shown as motif occurrence percentage for shared motifs in H3K27me3 (hot pink) and H3K36me2 (light green) (**H**); and H3K27me3 (hot pink) and H3K4me3 (light blue) (**I**); consensus sequences for representative motifs are shown for SPI1, KLF4, E2F6 and ZNF263 (**J**). (**K**) Analysis of gene expression levels is shown for EZH1 and EZH2 in control (grey box) and E2148* (light blue box) mutant neurons.

Functional enrichment analysis by GO categories across H3K27me3 differential peaks in gene encoding regions identified an overrepresentation of pathways involved in establishing the wiring of the brain (i.e. axonogenesis, axon guidance, voltage-gated channel activity, potassium channel activity, glutamatergic synapses, and neuronal projection) in *ASH1L* mutant neurons (**Fig.6E-G and Supplementary table S13**). Additional examination of the enhanced H3K27me3 peaks showed enrichment for WNT signaling, cell-cell adhesion, extracellular matrix, and growth cone dynamics (i.e. cell leading edge and focal adhesion) across different GO categories (**Supplementary Fig. S7D-F and Supplementary table S13**). The functional enrichment analysis of H3K27me3 differential and enhanced peaks inversely mirrors our findings on H3K36me2 and H3K4me3 (**Fig. 5**, **supplementary Fig. S6 and Supplementary Table S10**) suggesting counteracting epigenomic regulatory mechanisms that are driven by ASH1L during the early stages of human brain development.

To identify additional regulomes modulated by ASH1L, we conducted motif enrichment analysis (**Fig. 6H-J and Supplementary Table S14**) across H3K27me3 differential peaks and compared them to motifs associated with H3K36me2 and H3K4me3. Overlapping motifs between H3K36me2 and H3K27me3 were enriched for Stat2, ZNF263 and KFL4 (**Fig. 6H and 6J**), while motifs shared between H3K4me3 and H3K27me3 showed higher prevalence for SP1, KLF4 and ZNF263 (**Fig. 6I-J**). To pinpoint regions co-regulated by ASH1L and PRC2 we analyzed the overlapping differential peaks for H3K36me2 and H3K27me3, identifying the associated regulome (**Supplementary Fig. S8A**). These regions showed significant enrichment for SP1, SP2, KLF4, and E2F6 motifs (**Supplementary Fig. S8B**). Therefore, our findings indicate that alterations in ASH1L function disrupt the chromatin environment at multiple levels, converging on gene programs relevant to neuronal structure and function. This convergence is potentially driven by SP and KLF transcription factor families, which are essential regulators of brain development.

To determine whether the increase in genome-wide loci marked by H3K27me3 was associated with increased expression of the catalytic subunits of PRC2, namely EZH1, and EZH2, we examined their gene expression levels but found no significant changes (**Fig.6K**). To further interrogate whether the genome-wide increase in H3K27me3 differential peaks reflected higher H3K27me3 total levels in neurons, we conducted western blot analysis on chromatin fractions. We found that while variable there were no significant changes in global H3K27me3 levels (**Supplementary Fig.S8C**). To further interrogate if the lack of a significant increase in total levels of H3K27me3 could result from mechanisms to preserve cellular homeostasis, we measured protein levels for different PRC2 components. We found decreased protein levels for catalytic subunits EZH2 and EZH1, while SUZ12 and the non-catalytic subunit EED remained unaffected (**Supplementary Fig. S8D**). Importantly, this reduction in PRC2 subunit proteins is not due to decreased gene expression, as RNA-seq showed no significant changes in their mRNA levels (**Fig. 6K**). This evidence suggests that neurons with ASH1L dysfunction might try to compensate for the increased locus-specific H3K27me3 levels by regulating the PRC2 catalytic subunits at the protein level. However, these efforts may be insufficient to prevent transcriptional repression of critical neurodevelopmental pathways.

### Epigenetic strategies that promote transcriptional activation rescue defects in neurite outgrowth and morphogenesis associated with ASH1L dysfunction

We previously showed that inhibition of EZH2, the catalytic subunit of PRC2, rescues neuronal arborization defects caused by shRNA-mediated ASH1L knockdown in neurons derived from human embryonic stem cells ^40^. Similarly, through a separate mechanism of histone deacetylation inhibition that similarly results in transcriptional activation, the small molecule Vorinostat rescued ASD-like behaviors in ASH1L mutant mice ^28^. Therefore, harnessing different epigenetic mechanisms that modulate transcriptional activation could provide a potential rescue strategy for ASH1L-related cellular phenotypes. Here we focused on two FDA-approved small molecules used in cancer treatment – Tazemetostat, an EZH2 inhibitor ^67,68^ and Vorinostat, a histone deacetylase (HDAC) inhibitor ^27^ (**Fig. 7 and Supplementary Fig. S9**). After determining the optimal doses for human neurons through viability assays, (**Supplementary Fig. S9A-B**) control and *ASH1L* mutant neurons were treated for 3 days with 0.5 µM Tazemetostat and 0.1 µM Vorinostat and analyzed for changes in neuronal morphogenesis (**Fig. 7A-F**). Analysis of changes in gene expression for ASH1L showed that neither treatment altered ASH1L expression as levels remained significantly lower than the control in both mutant lines (**Supplementary Fig. S9E**). *ASH1L* mutant neurons showed a significant increase in total and mean neurite length after treatment with either Vorinostat or Tazemetostat, reaching levels comparable to the DMSO-treated controls (**Fig. 7A-C**). Analysis of the complexity index, a measure of neuronal arborization, showed that Vorinostat preferentially rescued the arbor complexity to levels significantly different from the untreated mutant neurons, while Tazemetostat showed a similar trend but was not significant (**Fig. 7D**). Sholl analysis confirmed increased arborization with both Tazemetostat and Vorinostat compared to the untreated E2148* mutant neurons (**Fig. 7E**), with a more significant effect associated with Vorinostat treatment to levels similar to the control neurons (**Fig. 7F**).

**Figure 7.**
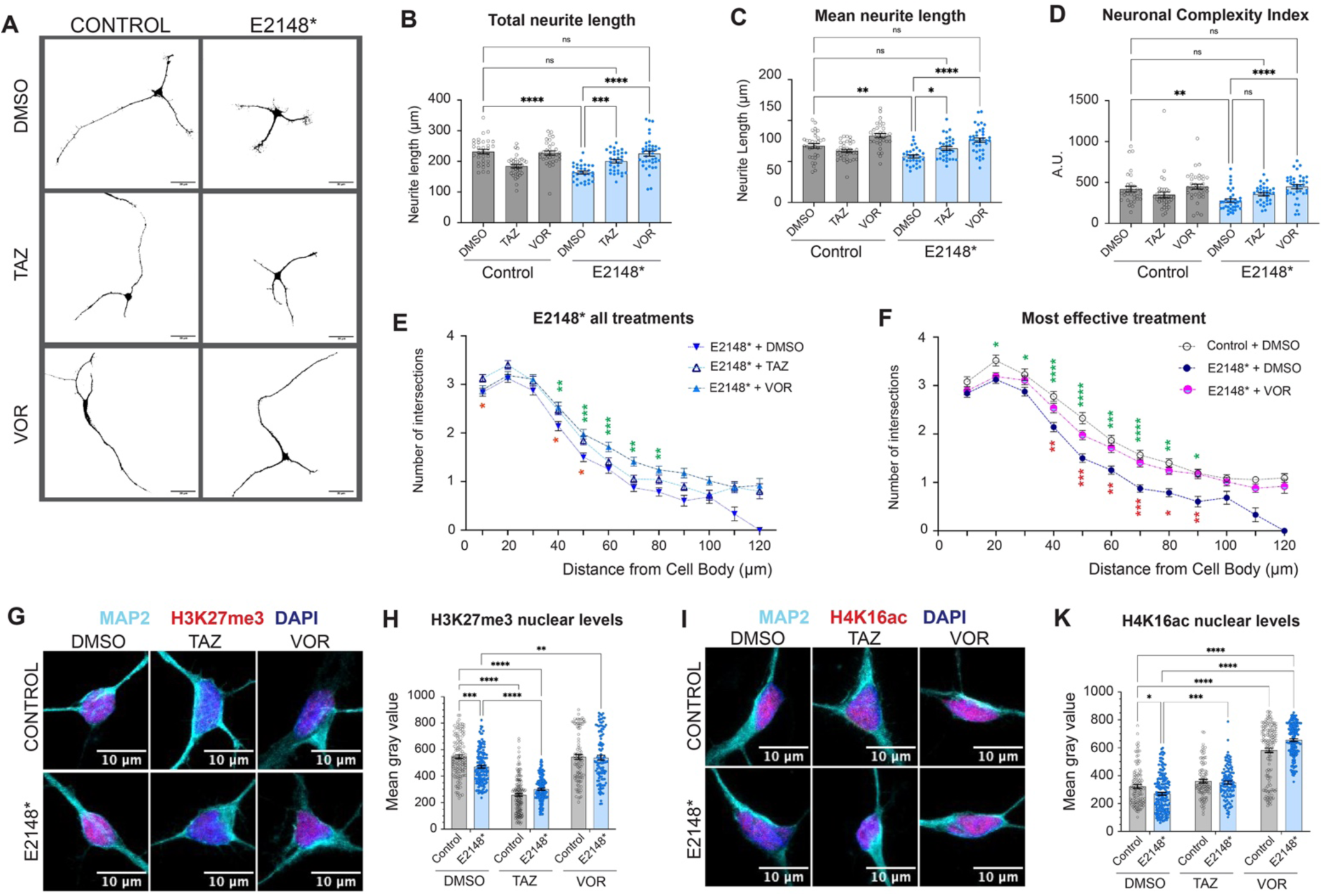
Tazemetostat and Vorinostat rescue neuronal arborization in ASH1L mutant neurons by targeting H3K27me3 and H4K16ac levels. **(A)** Representative images are shown for day 35 human neurons from control, and E2148* cultures treated for 3 days with DMSO, Tazemetostat (0.5µM) and Vorinostat (0.1µM). Neurons stained with MAP2 are shown in black and white for ease of viewing. Calibration bars represent 30µm. (**B-F**) Morphogenesis analysis is shown for at least 4 independent experiments (unless otherwise annotated) in which we measured at least 30 neurons per experiment for control (grey bar with open circles) and E2148* (light blue bars with solid dark blue circles) neurons treated with either DMSO, Tazemetostat (TAZ) or Vorinostat (VOR). Individual points represent the average of multiple independent experiments. (**B**) Total neurite length is shown as the mean (bar) with the average of individual measurements represented by the circles for: control + DMSO (n=93 neurons; 231.9 ± 7.27); control + TAZ (n=119; 184.4± 5.46); control + VOR (n=118 neurons; 227.3± 6.8); E2148* + DMSO (n=113 neurons; 163.3 ± 4.83); E2148* + TAZ (n=112; 200.5± 5.94); E2148* + VOR (n=129 neurons; 225.2± 8.38). (**C**) Mean neurite length is shown as the mean (bar) with the average of individual measurements represented by the circles for: control + DMSO (n=90 neurons; 69.86 ± 2.409); control + TAZ (n=117; 64.16± 2.04); control + VOR (n=118 neurons; 81.28± 2.75); E2148* + DMSO (n=112 neurons; 55.75 ± 1.88); E2148* + TAZ (n=111; 66.16 ± 2.47); E2148* + VOR (n=126 neurons; 76.02 ± 2.58). (**D**) Complexity index measurements were first analyzed using the “identify outliers” ROUT function in graph pad and are shown as the mean (bar) with the average of individual measurements represented by the circles for: control + DMSO (n=90 neurons; 390± 26.88); control + TAZ (n=111; 329.4 ± 22.40); control + VOR (n=116 neurons; 461.8 ± 25.45); E2148* + DMSO (n=116 neurons; 261.9 ± 18.80); E2148* + TAZ (n=105; 357.6 ± 20.83); E2148* + VOR (n=125 neurons; 458.3 ± 29.66). (**B-D**) Statistical analysis of grouped measurements was conducted using TWO-way ANOVA with Tukey’s test for multiple comparisons: * P < 0.04 ** P < 0.009, *** P < 0.0006, **** P < 0.0001. (**E**) Sholl analysis was used to measure neuronal arborization across three different treatments in the E2148* mutant neurons. The number intersections away from the cell soma were measured every 10µm and are shown for E2148* + DMSO (inverted dark blue triangles), E2148* + TAZ (open triangles), and E2148* + VOR (solid light blue triangles) neurons. Statistical analysis by TWO-way ANOVA with mixed model effects * P < 0.05, ** P < 0.009, and *** P = 0.0008. Green asterisk (E2148* +DMSO vs. E2148* + VOR), red asterisk (E2148* + DMSO vs. E2148* + TAZ). (**F**) Sholl analysis is shown to compare the most effective treatment (vorinostat) to the untreated control and E2148* mutant neurons. The number of intersections away from the cell soma were measured every 10µm and are shown for control+ DMSO (open gray circles), E2148* + DMSO (solid dark blue circles) and E2148* + VOR (half pink/light blue circles) neurons. Statistical analysis by TWO-way ANOVA with mixed model effects * P < 0.05, ** P < 0.005, *** P = 0.0005, and **** P < 0.0001. Green asterisk (E2148*+DMSO vs. Control + DMSO), red asterisk (E2148* + DMSO vs. E2148* + VOR). (**G-K**) Analysis of nuclear levels of H3K27me3 and H4K16ac in four independent experiments (unless otherwise indicated) across all treatments is shown for neurons at day 35 of neuronal induction. (**G**) Representative images of nuclear H3K27me3 (red) are shown for either DMSO (left column) or Tazemetostat (right column) treated control, or E2148* mutant neurons stained with MAP2 (cyan) and nuclei is stain with DAPI (blue). (**H**) Quantification of H3K27me3 nuclear levels measured by mean gray value is shown for all treatments. Measurements from at least 3 independent experiments with at least 30 neurons analyzed per experiment were analyzed as a group and are shown as the mean (bar) with the average of individual measurements represented by the circles for: control + DMSO (n=134 neurons; 547.4 ± 13.19); control + TAZ (n=154; 258.1 ± 10.83); control + VOR (n=101 neurons; 545.5 ± 19.07); E2148* + DMSO (n=141 neurons; 471.0 ± 10.87); E2148* + TAZ (n=140; 301.1 ± 8.03); E2148* + VOR (n=103 neurons; 539.4 ± 18.79). (**I**) Representative images of nuclear H4K16ac (red) are shown for either DMSO (left column) or Tazemetostat (right column) treated control, and E2148* mutant neurons stained with MAP2 (cyan) and nuclei is stain with DAPI (blue). (**J**) Quantification of H4K16ac nuclear levels measured by mean gray value is shown for all treatments. Measurements from at least 3 independent experiments with at least 30 neurons analyzed per experiment were analyzed as a group and are shown as the mean (bar) with the average of individual measurements represented by the circles for: control + DMSO (N= 4 experiments; n=114 neurons; 322.8 ± 12.85); control + TAZ (n=95; 360.5 ± 13.04); control + VOR (n=145 neurons; 581.6 ± 16.45); E2148* + DMSO (n=166 neurons; 270.3 ± 10.33); E2148* + TAZ (n=106; 350.4 ± 11.81); E2148* + VOR (n=158 neurons; 654.7 ± 9.49). (**H and J**) Statistical analysis of grouped measurements was conducted using TWO-way ANOVA with Tukey’s test for multiple comparisons: * P < 0.05, *** P < 0.005, *** P < 0.0005, **** P < 0.0001.

To validate the mechanisms of action for Tazemetostat and Vorinostat in the human neurons, we assessed nuclear levels of H3K27me3 and acetylation of lysine 16 on histone H4 (H4K16ac) through immunofluorescence (**Fig. 7G-K**). Focus on H4K16ac was based on previous findings that ASH1L knockdown in non-neuronal cells reduces H4K16ac levels at target genes ^69^. We observed a discrete but significant decrease in H3K27me3 levels in the mutants treated with DMSO compared to controls, while Tazemetostat treatment reduced nuclear H3K27me3 by nearly two-fold in both control and mutant neurons (**Fig. 7G-H**). Interestingly, Vorinostat treatment led to a modest but significant increase in H3K27me3 in the E2148* mutant neurons (**Fig. 7H**). H4K16ac nuclear levels showed a significant but discrete decrease in DMSO treated E2148* mutant neurons compared to DMSO treated control neurons (**Fig. 7I-K**). As expected, treatment with the HDAC inhibitor Vorinostat raised H4K16ac levels in mutant and control neurons (**Fig. 7K**) by at least 2.4-fold compared to their respective DMSO-treated controls. Unexpectedly, Tazemetostat also resulted in a significant but modest increase in nuclear H4K16ac levels in the mutant neurons (**Fig. 7K**). Taken together, these findings suggest that Tazemetostat-induced downregulation of H3K27me3 and Vorinostat-induced upregulation of H4K16ac primarily underlie the rescue of neuronal morphogenesis; however, the results also highlight crosstalk between these epigenetic rescue strategies within the neuronal chromatin landscape.

## DISCUSSION

Human iPSC-derived neurons provide a robust platform for studying cell autonomous phenotypes throughout early neuronal development ^70–72^. Here we harness the power of iPSC systems and CRISPR-CAS9 genome editing to establish an isogenic model of ASH1L disease to interrogate the effect of mutating the catalytic domain of ASH1L during neuronal development. The various clinical phenotypes associated with mutations in ASH1L highlight the importance of this chromatin regulator in human neuronal development ^6–17^. However, the precise cellular phenotypes and molecular signatures that underlie the pathogenesis of ASH1L-related disorders in human neurons have been largely understudied. Our work bridges this gap by providing an in-depth analysis of alterations in neuronal transcriptome, alternative splicing, and the chromatin environment in human neurons with disruption in the catalytic domain of ASH1L. Patients with ASH1L-disease present a multiple array of clinical presentations; however, the precise contributions of the genetic background or domain-specific mutations in ASH1L-related disorders is unknown. The present study using a male 11A iPSC ^33^ line and our previous work using a male H1 embryonic stem cell line ^40^ suggest that neuronal arborization defects associated with ASH1L dysfunction are independent of genetic background. Additionally, our previous and present work further suggests that the rescue of neuronal arborization by targeting the catalytic activity of PRC2 is also independent of genetic background.

We define a novel functional transcriptomic and epigenetic node that could alter neuronal structure and synaptic function development in response to deficits in ASH1L catalytic activity. At the transcriptome level, we observed a preference for the downregulation of longer genes that could result partly from the reduction in transcription rates observed in neurons with ASH1L dysfunction that showed lower incorporation of nuclear EU. Changes in transcription efficiency in long genes could result from disruptions in topology-associated domains due from genome-wide changes in the histone modification landscape^73^. Similarly, decreased transcriptional elongation caused by inhibition of Topoisomerase I (TOP I) was previously shown to reduce the expression of extremely long ASD genes ^46^. Similarly, Rett syndrome MECP2 was shown to repress the expression of long neuronal genes by modulating Topoisomerase II (TOP-II) function ^47^. TOPI/II regulation of long gene expression could be a shared mechanism underlying ASD pathogenesis. Therefore, an alternate mechanism through which ASH1L regulates neuronal long gene expression could be mediated by TOP I/II function, and this might be a shared mechanism of pathogenicity between ASH1L-related disorders and Rett syndrome.

Our work implicates that discrete changes in alternative splicing could be contributing to the neuronal defects associated with ASH1L dysfunction. In particular, we highlight splicing alterations in *SMARCA4*, which is an important neurodevelopmental gene that regulates chromatin remodeling. The fact that *SMARCA4* is not differentially expressed in our RNAseq dataset stresses the importance of considering disruption in alternative splicing in relation to defects in chromatin regulators like ASH1L, which is often overlooked as a mechanism of disease pathogenesis in relation to ASD. Based on our biochemical analysis of MRG15, EU incorporation studies, and analysis of H4K16ac nuclear levels, we posit two alternative mechanisms that could impact alternative splicing: (**1**) reduced levels of chromatin-bound MRG15 could lead to decreased recruitment of splicing factors to the pre-mRNA; or (**2**) alterations in histone acetylation could change RNA polymerase elongation rates and contribute to defects in alternative splicing ^74^. Future studies from our group will dissect the contribution of these mechanisms in the control of alternative splicing by ASH1L.

Next, we identified synaptic gene networks regulated by ASH1L in human neurons. We expect that similar to the in vivo studies in which ASH1L mutant mice show defects in synaptic function ^75^ ASH1L will also regulate synaptic activity in humans. Further, because the chromatin profile of H3K27me3 is also altered across synaptic structure and function loci, we posit that potential defects in neuronal activity will be rescued with the same strategies we used to rescue the neuronal morphogenesis phenotypes.

With respect to the small molecule rescue approaches, the discrete differences in the treatment response depending on the type of metrics used (complexity index vs. Sholl analysis) suggest the future need for individualized therapeutic strategies and the need to test these strategies in patient-derived iPSC systems using multiple metrics. These models will consider the contribution of the genetic background to the cellular and molecular phenotypes. We are aware of the limitations of using broad dysregulation of epigenetic mechanisms as a therapeutic strategy to treat ASH1L-related neurological disorders, and this emphasizes the need to identify and develop additional targeted therapeutic strategies. Additionally, our work highlights the crosstalk across different epigenetic mechanisms and how they could influence different rescue strategies. We showed that Vorinostat and Tazemetostat altered not only their intended histone mark but in the case of Tazemetostat there was an additive effect as it decreased H3K27me3 levels and modestly increased H4K16ac levels in the mutant neurons. In contrast, Vorinostat almost doubled H4K16ac levels and modestly increased H3K27me3 in neurons with defects in ASH1L’s catalytic domain. Therefore, the interplay between different histone modifications and how they could affect multiple transcriptional programs should also be considered when targeting different epigenetic mechanisms as rescue strategies.

Finally, we identify the SP and KLF families of transcription factors as novel regulons modulated by the ASH1L/PRC2 axis in human neurons. Members of the SP and KLF families have been associated with different aspects of neuronal development including neurogenesis, neuronal morphogenesis and synaptic function ^63–66^ . These cellular processes are enriched among the epigenomic and transcriptomic programs modulated by ASH1L in human neurons, and further emphasize the importance of ASH1L in human brain development.

## ONLINE METHODS

### Stem cell culture

Human 11A induced pluripotent stem cells (iPSCs) ^33^ obtained from a neurotypical male individual were a gift from the Kevin Eggan laboratory and were obtained from the Harvard Stem Cell Institute (Boston, MA) and were used to introduce pathogenic variants in ASH1L. Isogenic control and mutant iPSCs colonies were inspected daily to ensure there was no spontaneous differentiation and were maintained in mTeSR™ Plus basal medium supplemented with 5X supplement (STEMCELL Technologies, #100-0276) on Matrigel (Corning #354277) coated plates as previously described ^36^. Cells were fed daily and were passaged every 3-4 days at a 1:6 ratio using ReLeSR™ (STEMCELL Technologies, #100-0483). Wide-bore micropipette tips (Thermo Scientific #2079G) were used to prevent the formation of single-cell suspensions. To minimize chromosomal abnormalities, iPSCs were maintained only up to passage 50.

### CRISPR/CAS9 genome editing

We used CRISPR/CAS9 genome editing to introduce mutations in the catalytic domain of ASH1L ^7,30,31^. We used the previously established iPSC line -11A, generated from a healthy, neurotypical male donor ^33,34^, to introduce a nonsense pathogenic mutation using the CRISPR/CAS9 RNP system ^35,36^. Clones were screened for heterozygous mutations by Sanger sequencing. Genomic DNA from mutant clones was sequenced to identify the mutations introduced into ASH1L.

### Generation of forebrain cortical excitatory neurons from human iPSCs

11A control and ASH1L mutant iPSCs were differentiated into forebrain cortical excitatory neurons as described previously ^36,42^. Briefly, human iPSC colonies were dissociated with Gentle Cell Dissociation Reagent (STEMCELL Technologies, #100-0485). Dissociated iPSCs were resuspended in mTeSR™ Plus medium (STEMCELL Technologies, #100-0276) supplemented with ROCK inhibitor Y27632 (STEMCELL Technologies, #72304) and plated at 1x 10^6^ cells per well into 12-well tissue culture plates coated with Matrigel (Corning #354277). After ensuring the complete formation of a monolayer on the next day, cell media was replaced with neural induction medium (NIM) made with 1:1 DMEM/F-12 GlutaMAX (Thermo Fisher Scientific, #10565018) and Neurobasal media (Thermo Fisher Scientific, #21103049) containing: N2 supplement (Thermo Fisher Scientific #17502-048), B27 supplement (Thermo Fisher Scientific, #17504044), 5µg/ml insulin (Sigma-Aldrich, #I0526-5ML), 1mM L-glutamine (Thermo Fisher Scientific, #25-030-081), 100 μM nonessential amino acids solution (Thermo Fisher Scientific, #11-140-050), 100µM β-mercaptoethanol (Sigma-Aldrich, #M6250-100ML), 50U/ml penicillin-streptomycin (Thermo Fisher Scientific, #15-140-122), 1µM Dorsomorphin (STEMCELL Technologies,#72102) and 10µM SB431542 (STEMCELL Technologies, #72234). Between days 12 to 16, the presence of neuronal rosettes composed of neuronal progenitors indicated the success of the neuronal induction. Neuronal progenitors were expanded in NIM with the addition of 10µM recombinant FGF2 (PeproTech, #100-18C) from days 17 to 21. After day 21, neurons were fed with half media changes every other day. Neurons were allowed to mature in neural maintenance medium (NMM), containing all the components of NIM except for Dorsomorphin and SB431542. For subsequent experiments neurons were seeded on either 8-well chamber slides or multi-well tissue culture plates coated with 20µg/ml laminin (Sigma-Aldrich, #L2020-1MG) and poly-ornithine (Sigma-Aldrich, #P4957-50ML).

### Quantitative PCR for Gene Expression Analysis

To measure changes in gene expression in either neurons or iPSCs, RNA was isolated from control and mutant lines using QIAGEN RNeasy Mini kit (Qiagen #74104). RNA concentration was measured using Invitrogen Qubit 4 Fluorometer using the RNA Broad Range quantification assay (Qubit #Q10210). cDNA synthesis was performed using Superscript IV First-Strand cDNA Synthesis Kit (Thermo Scientific #18091050). Quantitative PCR (qPCR) was carried out using TaqMan Gene Expression Assays (Thermo Scientific # 4331182, **Supplemetary Table S2**) targeting the following genes: OCT4, NANOG, SOX2, ASH1L, MAP2, RLP13A. Gene expression levels were normalized using the ΔΔCq method ^76^, with fold changes calculated relative to reference gene RPL13A for iPSCs or MAP2 for neurons. Each experiment was performed with four independent biological replicates.

### EU Incorporation Assay

Click-iT® RNA Alexa Fluor® 594 Imaging Kit (Invitrogen, C10330) was used for the EU (5-ethynyl Uridine) incorporation assay. Briefly, the stock solutions were prepared as per the manufacturer’s instructions. After the treatment, a working solution of 100 mM stock of EU was added to the well for a final concentration of 100 μM in each well, followed by incubation for 3 hours at 37 °C. The media was then removed, and the wells were washed twice with 1XPBS (Thermo Scientific, #70011044) before fixing them using 4% paraformaldehyde (Electron Microscopy Sciences, #15710-S) for 15 minutes at room temperature. Cells were permeabilized and washed with 1XPBS, followed with a 30-minute incubation in the dark with freshly prepared 1X Click-iT solution and Click-iT cocktail. After 30 minutes, cells were washed with Click-iT rinse buffer. Primary antibody for MAP2 (Aves lab, #MAP) was added and cells were incubated overnight at 4°C, followed by secondary antibody incubation for 1 hour at room temperature using Alexa Fluor 647 (Thermo Scientific, #A-21449) at a dilution of 1:1000. Finally, the coverslips were mounted using mounting medium, Prolong Gold Antifade (Invitrogen, #P36941) and imaged with FV3000 Olympus Confocal Microscope using oil immersion 60X objective. The quantification for total EU in the nucleus was done using ImageJ (Version 1.53c), with matched imaging parameters for exposure, gain, and offset (see image analysis section).

### Western blotting

For western blot analysis, neurons at day 41 were washed twice with cold 1X PBS and lysed in cell lysis buffer (20 mM Tris-HCl (pH 7.5), 150 mM NaCl, 1 mM Na_2_EDTA, 1 mM EGTA, 1% Triton, 2.5 mM sodium pyrophosphate, 1 mM beta-glycerophosphate, 1 mM Na_3_VO_4_, 1 µg/ml leupeptin supplemented with protease inhibitor). The lysates were centrifuged at 12,000 *g* for 15 min at 4 °C to separate the chromatin-bound and soluble fractions. Lysates were quantified using Pierce BCA Protein Assay kit (Thermo Scientific #23227), and equal amounts of protein were loaded onto a 4 to 12% gradient gel (NuPAGE Invitrogen). Protein was transferred from the gel to 0.2 μm NC membranes at 25 V for 10 min using transfer stacks (iBlot2 Invitrogen) and blocked with LiCor TBS blocking buffer (#927-60003) for 1 hour at room temperature before application of primary antibodies. Primary and secondary antibodies were incubated overnight at 4 °C and for 1 hour at room temperature, respectively. Membranes were incubated with primary antibodies using the following dilutions: MRG15 (Proteintech, #55257-1-AP 1:1000), EZH2 (Cell Signaling Technology, #5246S 1:1000), EZH1 (Cell Signaling Technology, #42088S 1:1000), SUZ12 (Cell Signaling Technology, #3737S 1:1000), EED (Cell Signaling Technology, #85322S 1:1000), Histone H3 (Cell Signaling Technology, #14269S 1:1000), H3K4me3 (Cell Signaling Technology, #9751S 1:500), H3K36me2 (Cell Signaling Technology, #2901S 1:1000), H3K36me3 (Cell Signaling Technology, #4909S 1:500), H3K27me3 (Cell Signaling Technology, #9733S 1:1000), and GAPDH (Cell Signaling Technology, #97166S 1:1000) prepared in LiCor Intercept T20 (TBS) Antibody Diluent (#927-65003). Blots were washed 1X TBST (10 mM Tris-HCl pH 8.0, 150 mM NaCl, 0.01% Tween-20) after primary antibody incubation. Fluorescent secondary antibodies, specifically, anti-rabbit (Thermo Scientific #A32729) and anti-mouse (Thermo Scientific #A32735) were used at 1:5000 dilutions. The blots were imaged using LiCor Odyssey CLX scanner. Quantification of western blots was done using ImageJ (Version 1.53c).

### Immunocytochemistry

At day 32 of neuronal induction, iPSC-derived human cortical neurons were plated (30,000 cells/well) in 8-well chamber slides (CELLTREAT, #229168) coated with 0.01% poly-L-ornithine and 10µg/ml laminin. For each experiment four independent neuronal inductions were used. At day 35 of neuronal induction cells were briefly rinsed with 1X PBS (Thermo Scientific, #70011044), fixed with 4% paraformaldehyde (Thermo Scientific, 043368.9M) for 15 minutes (min) at room temperature, washed three times for 5 min each with 1XPBS, permeabilized with 0.25% Triton X-100 for 10 min at room temperature, washed three times for 5 min each with room temperature 1X PBS, and blocked with 10% goat serum solution (Thermo Scientific, #50062Z) for one hour at room temperature. After blocking, primary antibodies against either MAP2 (Abcam #36592 1:1250), H3K27me3 (Active Motif #39155 1:300) and H4K16ac (Cell Signaling Technology #13534 1:1200) were diluted in 2% goat serum, 0.2% Triton X-100^36^ were applied to cells and incubated overnight at 4°C. Following primary antibody treatment, cells were washed three times with 1XPBS for 5 minutes each time, and then incubated for 1 hour at room temperature with secondary antibodies diluted in 1XPBS with 2% goat serum (AlexaFluor-594 anti-mouse Thermo Fisher Scientific, #A11032, Alexa Fluor-647 anti-rabbit Thermo Fisher Scientific # A-21245, and Alexa Fluor-488 anti-chicken Thermo Fisher Scientific, # A32931, all diluted at 1:1000). Cells were washed three times with 1XPBS, and slides were mounted with ProLong Glass Antifade Mountant with NucBlue Stain (Thermo Scientific #P36985) under 1 mm rectangular coverslips (Fisher Scientific, #12-548-5P).

### Image acquisition and analysis

#### Imaging

Neuronal images were captured using an FV3000 Olympus Confocal Microscope equipped with a 60X or a 20X objective lens. Images taken with a 20X magnification lens were enlarged using the digital zoom for an additional 2X to 3.95X, optimized to prevent loss of resolution. Fluorescent images of 30 to 35 random cells were taken for at least four independent experiments.

#### Morphological Analysis

Morphological analysis was performed by experimenters blinded to genotype and experimental conditions using MBF Bioscience Neurolucida 360 and Neurolucida Explorer software. Neurites were traced from the cell body, and summary data for each cell were recorded, including total neurite length, mean neurite length, number of neurites, and complexity index (calculated as the total number of branches from neurite terminal endings to the cell body multiplied by the average neurite length). Additionally, spatial Sholl analysis was conducted to assess neuronal arborization, defined as the number of intersections between neurites and concentric circles at increasing 10 µm intervals from the cell body. A minimum of three independent experiments were conducted per condition. Neurite summary data were compared using the group function in GraphPad Prism.

#### Colocalization analysis

Protein colocalization in neuronal cells was assessed for two different fluorophores by generating a maximum intensity z-projection from image stacks. Regions of interest (ROIs) were created around individual cells using the select tool in Fiji ImageJ, and individual masks were generated to exclude external signals. The ROI was duplicated in split color channels—red (H3K27me3 and H4K16ac) and blue (DAPI). Pixel colocalization was measured using Pearson’s correlation coefficient with the Fiji ImageJ plugin JACoP^77^ and Coloc2^78^. Pearson’s correlation coefficient was chosen for its robustness to differences in pixel intensity, ensuring that biological changes in signal intensity across experimental groups are accounted for and preventing thresholding bias.

#### Protein levels intensity analysis

Protein level intensity was quantified by calculating the mean gray pixel value in Fiji ImageJ. ROIs were generated around individual cells as described previously, and color channels were split according to the protein of interest. Pixel intensity values were thresholded uniformly across all images to ensure consistency in the analyzed areas and to exclude background noise. The software calculated the mean gray value, representing the average pixel intensity within the selected area. These measurements were grouped per independent neuronal induction and were compared for analysis.

### Cell Viability and Neuronal Morphology Assays in response to small molecule treatments

For cell viability assays, neurons at day 32 of differentiation were seeded at 10,000 cells per well in 96-well plates. Viability was assessed 72 hours after treatment with Vorinostat (Selleck Chemicals, #S1047) and Tazemetostat (Selleck Chemicals, #S7128) at different concentrations using the 3-[4,5-dimethylthiazole-2-yl]-2,5-diphenyltetrazolium bromide (MTT) (Invitrogen, # M6494) assay. Following the 72-hour treatment, neurons were incubated with MTT (0.5 mg/mL) for 2 hours. The resulting insoluble purple formazan, produced by reduction of MTT by NAD(P)H-dependent oxidoreductases present in neurons with viable mitochondria, was solubilized in dimethyl sulfoxide (DMSO) at room temperature with agitation, and protected from light. The percentage of MTT reduction was determined by measuring the absorbance at 570 nm using a BioTek Cytation 5 plate reader. Results are presented as a percentage relative to control wells treated with DMSO. For neuronal morphological analysis, neurons at day 32 were plated onto 8 well chamber slides as previously described and treated with either DMSO (0.1%), 0.5 μM of Tazemetostat, or 0.1 μM of Vorinostat diluted in Neural Maintenance Medium (NMM). After 72 hours, neurons were fixed, processed for immunocytochemistry and imaged using a confocal microscope as previously described.

### Library preparation and sequencing

RNA was extracted using the RNeasy Micro Kit (Qiagen, #74004) following the manufacturer’s instructions. For each sample, RNA quality was analyzed using Agilent 2200 Tape Station system. cDNA libraries were prepared with the Collibri™ Stranded RNA Library Prep Kit for Illumina™ Systems (Invitrogen, #A38996096). Sequencing was performed using 150-bp reads using paired-end chemistry on an Illumina NovaSeq, to achieve a depth of 100 million reads.

### RNA-seq data processing and analysis

We assessed the quality of raw sequence data using FastQC (version 0.11.9) and TrimGalore (version 0.6.6). Raw reads were mapped to the Ensembl reference genome 104 (GRCh38) using the transcript-level quantifier Salmon (version 1.5.2) in mapping-based mode. Count matrices were generated from transcript-level quantification files using the tximport package (version 1.30.2). All RNAseq data is publicly available through GEO database associated with study number **GSE282613**. Differentially expressed genes (DEGs) were identified with DESeq2 (version 1.32.0) ^79^, considering genes with an adjusted p-value < 0.05 and a fold change ≥ |1.5| as statistically significant (**Supplementary Table S3**). Pathway enrichment analysis (GO, KEGG, DisGeNET) was performed using the EnrichR (version 3.2) and clusterProfiler (version 4.10.1) package in R. ASD associated genes used for **Fig. 4** were downloaded from SFARI database ^80^.

### Differential alternative mRNA splicing and transcript usage analysis

Isoform switching between mutants and controls was analyzed using the DEXSeq method implemented in the IsoformSwitchAnalyzeR package (version 2.2.0) ^81^ in R. This approach identifies bins with differential isoform switching between conditions, quantified as the difference in isoform fraction (dIF), calculated as IF_mutant - IF_control. The dIF values are effect size measurements, analogous to fold changes in traditional gene/isoform expression analysis. Significant isoform switching was defined by a dIF > 0.05 and a false discovery rate (FDR) < 0.05. Differential gene expression was determined for genes with a log2 fold change > 2 and a Q value < 0.05. Subsequent analysis included predictions of coding potential using CPC2 ^82^, protein domains using Pfam ^83^, signal peptide using SignalP-5.0 ^84^, intrinsically disordered regions (IDR) with IUPred2A ^85^ and topology using DeepTMHMM ^86^. Alternative splicing including exon skipping, intron retention, non-sense mediated decay status mutually exclusive exons were annotated throughout the analysis.

### Cleavage under Targets and Tagmentation (CUT&Tag) Assay

Genome-wide analysis of post-translational histone modifications, including H3K36me2, H3K27me3, and H3K4me3, was performed using the CUT&Tag kit (Active Motif #53160) following the manufacturer’s protocol. Human neurons, harvested at day 35 of neuronal induction, were used for this analysis. Briefly, 400,000 neurons were gently dissociated using Gentle Cell Dissociation Reagent (STEMCELL Technologies #100-0485) and incubated overnight with Concanavalin A beads and antibodies against H3K36me2 (Active Motif #39255), H3K27me3 (Active Motif #39155), H3K4me3 (Active Motif #39159), or a control rabbit IgG antibody (Cell Signaling Technology #2729S) at a concentration of 1 µg per reaction. Three biological replicates (defined as independent neuronal induction) were prepared per genotype for each antibody. Following incubation with the secondary anti-rabbit antibody (1:100), cells were washed, and tagmentation was conducted at 37°C using Protein A-Tn5. The tagmentation reaction was stopped by adding EDTA, SDS, and proteinase K, followed by DNA extraction, ethanol purification, and PCR amplification with barcoding, as per the Active Motif CUT&Tag protocol (#53160). DNA libraries were purified using SPRI beads (Beckman Coulter) and subsequently quantified and sequenced on an Illumina NovaSeq, generating close to 8 million 150 bp paired-end reads per sample.

### CUT&Tag Sequencing Analysis

Sequencing reads were aligned using to human genome (hg38) using Bowtie2 ^87^ with the following parameters: *--end-to-end --very-sensitive --no-mixed --no-discordant --phred33 -I 10 -X 700,* optimized for mapping insert sizes between 10-700 bps. The average overall alignment rate was 90%. Only high-quality reads (mapping quality ≥ 2) that were properly paired were retained for further analysis. Peak calling was performed using the MACS (version 2.1.0) ^88^ with a q value cutoff of 0.1, comparing each histone modification to the IgG controls. SAM files were generated, converted to BAM files, and biological replicates were merged using SAMtools ^89^. To represent the average distribution across each histone modification and condition. Heatmaps were created using DeepTools ^90^ from BigWig files, which were produced from the BAM filed using Bedtools ^91^. Differential peak analysis was conducted using the DiffBind package (version 3.12.0) ^92^ in R. Peaks with FDR < 0.05 were considered significant and were annotated using the Annotatr package (version 1.28.0) ^93^ in R to identify associated genomic regions. The distribution of histone modification sites relative to TSS was visualized using ChIPseeker (version 1.38.0)^94^ and rtracklayer (version 1.62.0) ^95^ with the hg38 genome assembly. The Integrative Genomics Viewer ^96^ (IGV version 2.16.2) was used to visualize the differential peaks across significantly different regions from the BigWig files for each condition.

#### Motif and gene regulatory region analysis

DNA sequences for differential peaks were extracted from the respective Bed files using Bedtools and rtracklayer (version 1.62.0) ^95^ with the hg38 genome assembly. The Integrative Genomics Viewer ^96^ (IGV version 2.16.2) was used to visualize the differential peaks across significantly different regions form the BigWig files for each condition. These sequences were analyzed for known motifs using the HOCOMOCO v12 CORE motif database using analysis of Motif Enrichment ^97^ (AME version 5.5.6) tool under default settings. Cis-regulatory regions were further predicted using the Genomic Regions Enrichment of Annotations Tool (GREAT) ^98^ for the Bed files containing the differential peaks, using default settings. All CUT&TAG seq data can be found under GEO accession numbers: **GSE282787, GSE282788, GSE282789.**

#### Pathway enrichment

Pathway enrichment analysis was performed for the gene lists associated with differential peaks using pathfindR^99^ (version 2.3.1) and ShinyGO (version 0.80) ^100^ for the Gene Ontology -Biological Process and Cellular Component gene sets.

## Supporting information

Jhanji_etal_Supplementary material

Jhanji_etal_Supplementary Tables

## ACKNOWLEDGEMENTS

We thank members of the Lizarraga laboratory (Dr. Elisa York) for critical reading of the manuscript. The research reported in this publication was supported by the National Institute of Mental Health of the National Institutes of Health under Awards Numbers R01MH127081 and R21 MH136643-01 to SBL and JSL. The content is solely the responsibility of the authors and does not necessarily represent the official views of the National Institutes of Health. This work was supported by a grant from Eagles Autism Foundation (Transcriptional mechanisms in ASH1L-related disorders, SBL and JSL). Illustrations were made with Biorender.

## AUTHORS CONTRIBUTIONS

SBL conceived the study, oversaw the experimental design and data analysis, and wrote the manuscript; MJ conducted all the biochemical studies, analyzed the transcriptomic and epigenomic studies, conducted the EU incorporation studies; contributed to the analysis of the arborization rescue studies, contributed to the overall data analysis, and wrote the manuscript; CL conducted the RNA seq and CUT&TAG experiments; JW conducted and analyzed the neuronal arborization and rescue experiments and gene expression analysis, SB provided pipelines and advice for analysis of epigenomic and transcriptomic datasets; CK, AG, and KV contributed to the biochemical studies, AG and KV contributed to iPSC gene expression analysis, imaging analysis; FDR contributed to the original analysis of pluripotency and neuronal arborization studies; BY contributed to the RNAseq analysis; KA contributed to sequencing analysis of the mutations. All authors contributed to the writing and editing of the manuscript.

