## Supplementary material for "Dynamic Regulation OF The Chromatin Environment By Ash1L Modulates Human Neuronal Structure And Function": Jhanji_etal_Supplementary material

Megha Jhanji [1,2], Joseph A. Ward [1,2*], Calvin S. Leung [1,2*], Colleen L. Krall [1,2], Foster D. Ritchie [3], Alexis Guevara[1,2], Kai Vestergaard[1,2], Brian Yoon [1,2], Krishna Amin[1,2,4], Stefano Berto [5], Judy Liu [1,2], and Sofia B. Lizarraga[1,2&]

[1] Department of Molecular Biology, Cell Biology and Biochemistry, Brown University, Providence, RI

[2] Center for Translational Neuroscience, Carney Brain Institute, Brown University, Providence, RI

[3] Department of Biological Sciences, Center for Childhood Neurotherapeutics, University of South Carolina, Columbia, SC

[4] Neuroscience graduate program, Brown University, Providence, RI

[5] Department of Neuroscience, Medical University of South Carolina, Charleston, SC

[*] Denotes equal contribution

**SUPPLEMENTARY MATERIAL**

Supplementary figures and figure legends as well as tables and /or table legends that are relevant to the main figures of the manuscript are presented.

**SUPPLEMENTARY FIGURES AND FIGURE LEGENDS**

**Supplementary Figure S1 relevant to Main Figure 1**

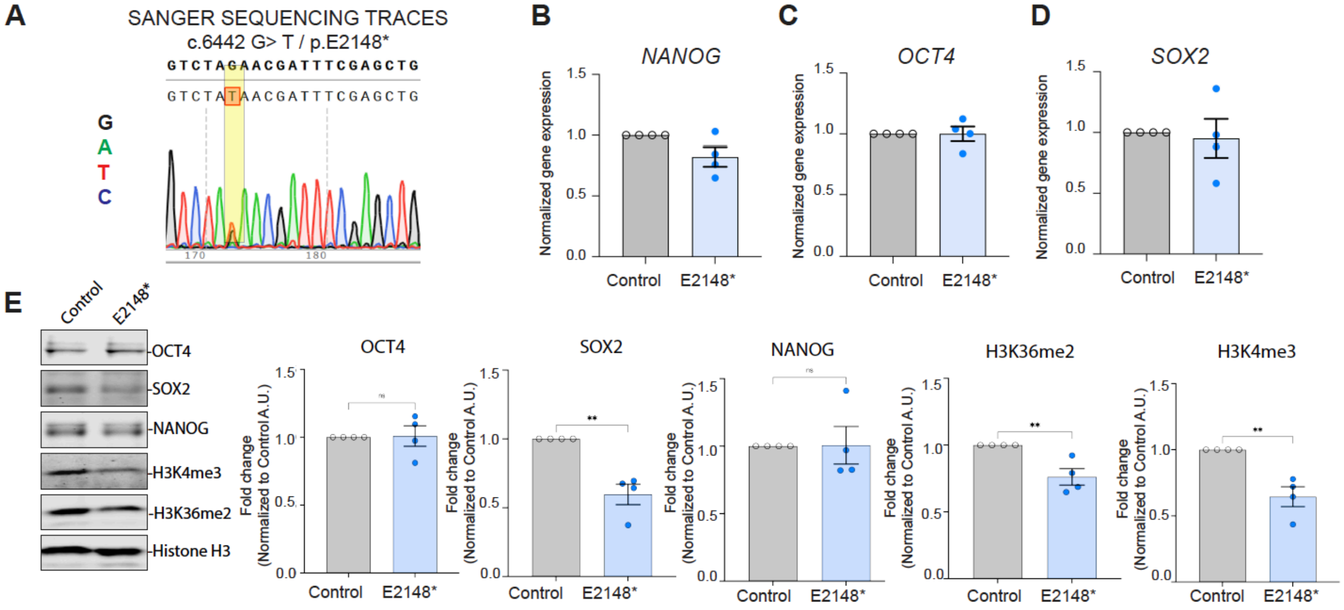


**Supplementary Figure S1. Characterization of iPSC pluripotency for ASH1L mutants. (A)** Sequencing traces showing the successful genome editing of pathogenic E2148* variant in ASH1L, from 11A male iPSCs as the parent line. **(B-D)** Characterization of pluripotency shows mRNA expression of *NANOG* (**B**), *OCT4* (**C**), and *SOX2* (**D**) in control, and E2148* iPSCs across 4 independent experiments (n=4). (**E**) Representative western blots and quantification show protein levels for pluripotency markers (OCT4, NANOG, SOX2) and histone methylation (H3K36me2 and H3K4me3) for control and E2148* mutant iPSCs (n=4). Error bars indicate standard error of the mean. Statistical analysis was performed using unpaired t-tests. Significant P- values are indicated as: *P<0.05, **P<0.01.

**Supplementary Figure S2 relevant to Main Figure 2**

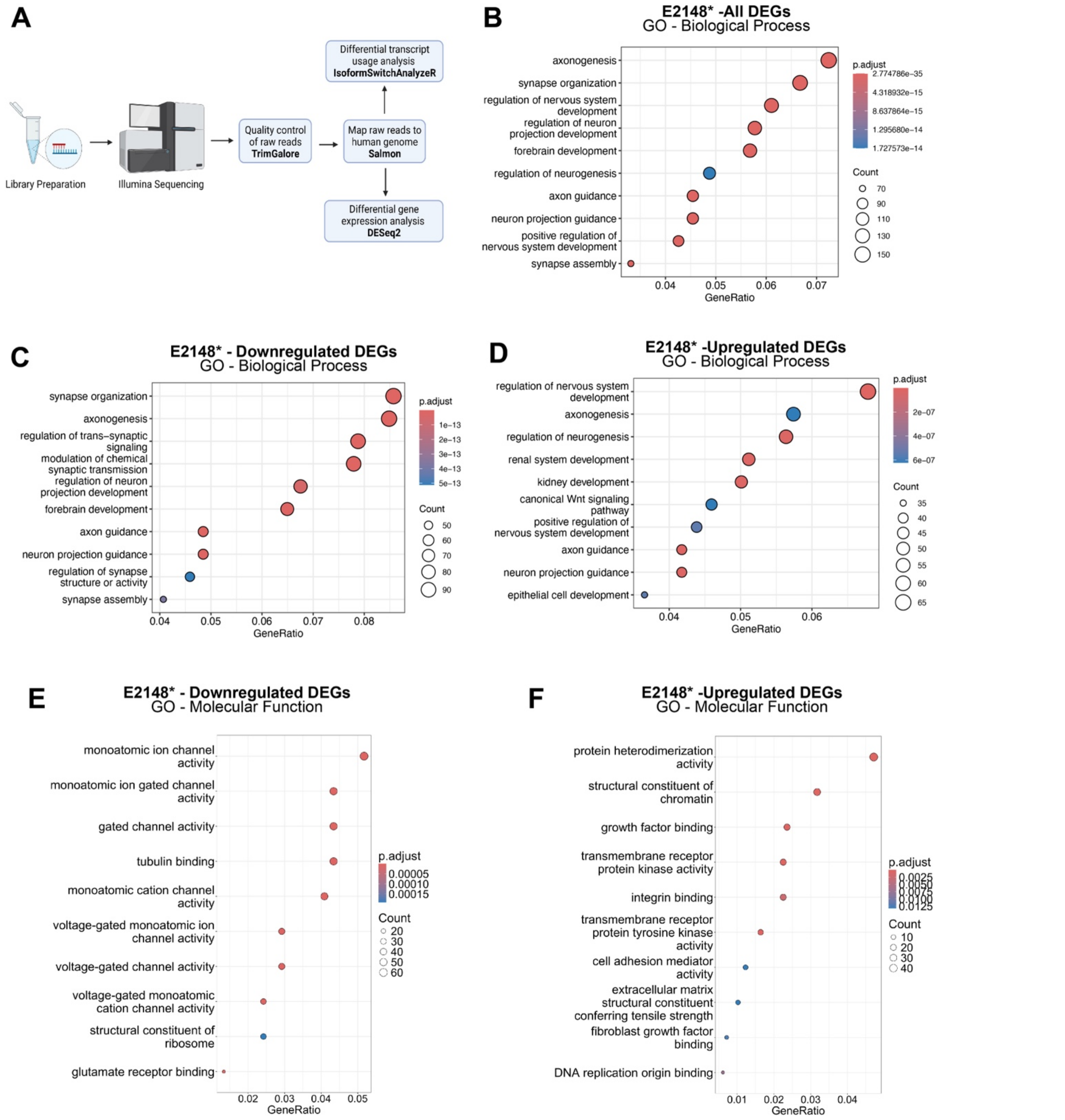


**Supplementary Figure S2. Transcriptome studies on ASH1L mutant neurons.** (**A**) Bioinformatics pipeline used for differential gene expression analysis and differential isoform analysis. (**B-D**) Gene ontology (GO) enrichment analysis for biological processes for DEGs E2148* mutant neurons (**B**), with downregulated (**C**) and upregulated DEGs (**D**). Circle size represents the number of DEGs in that category and the color represents the adjusted P value. (**E-F**) Gene ontology (GO) enrichment analysis for molecular function for upregulated (**E**), and downregulated DEGs (**F**). Circle size represents the number of DEGs in that category and the color represents the adjusted P value.

**Supplementary Figure S3 relevant to main figure 3**

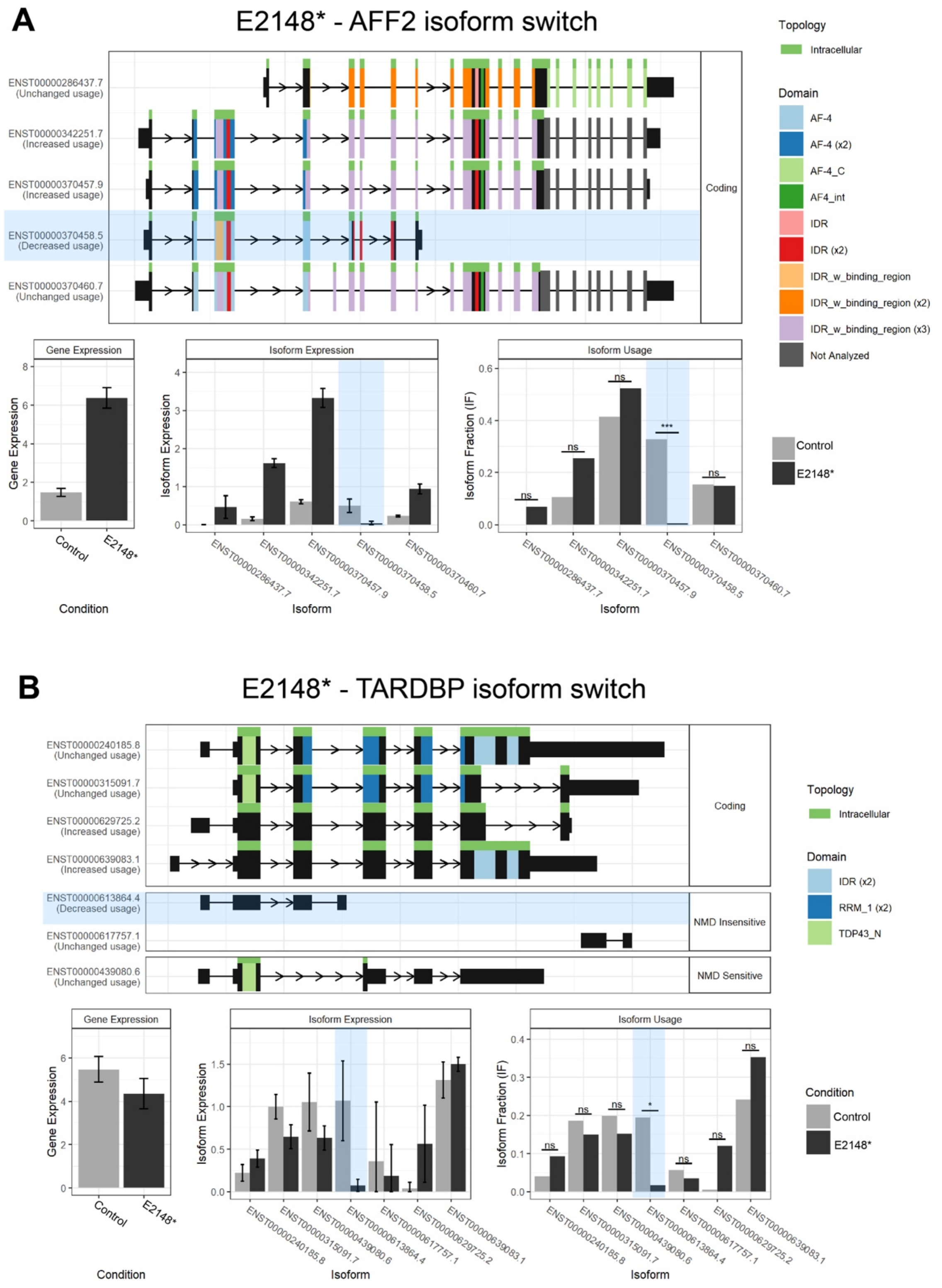


**Supplementary Figure S3. Differential isoform usage in E2148* human neurons.** (**A**) Graph shows all the coding and non-coding isoforms for AFF2. Analysis of isoform usage for AFF2 is shown for in control (grey) and mutant neurons (black) with the most significant changed isoform highlighted in blue. (**B**) Analysis of TARDBP gene and isoform expression in control (grey) and E2148* mutant neurons (black) with the most significant changed isoform highlighted in blue.

**Supplementary Figure S4 relevant to Main Figure 4**

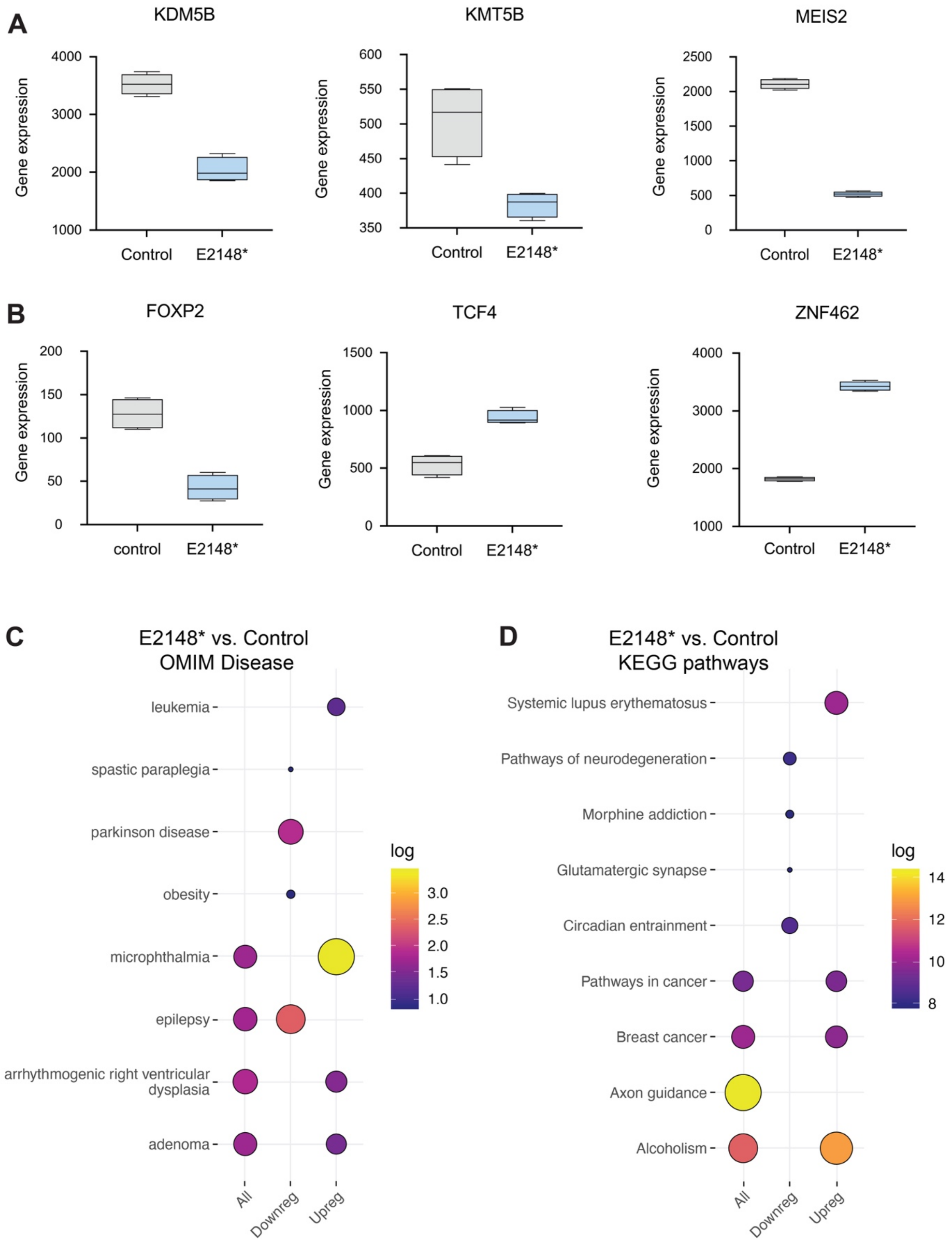


**Supplementary Figure S4. Differential gene expression of protein coding genes.** (**A-B**) Gene expression analysis by RNAseq of significant DEGs that are ASD-risk genes showing downregulated genes (**A**) and upregulated genes (**B**) for E2148* neurons vs control neurons. (**C**) Gene enrichment analysis for DEGs in OMIM disease gene sets. (**D**) Gene enrichment analysis is shown for DEGs in E2148* vs. control for KEGG pathways.

**Supplementary Figure S5 relevant to main figure 5**

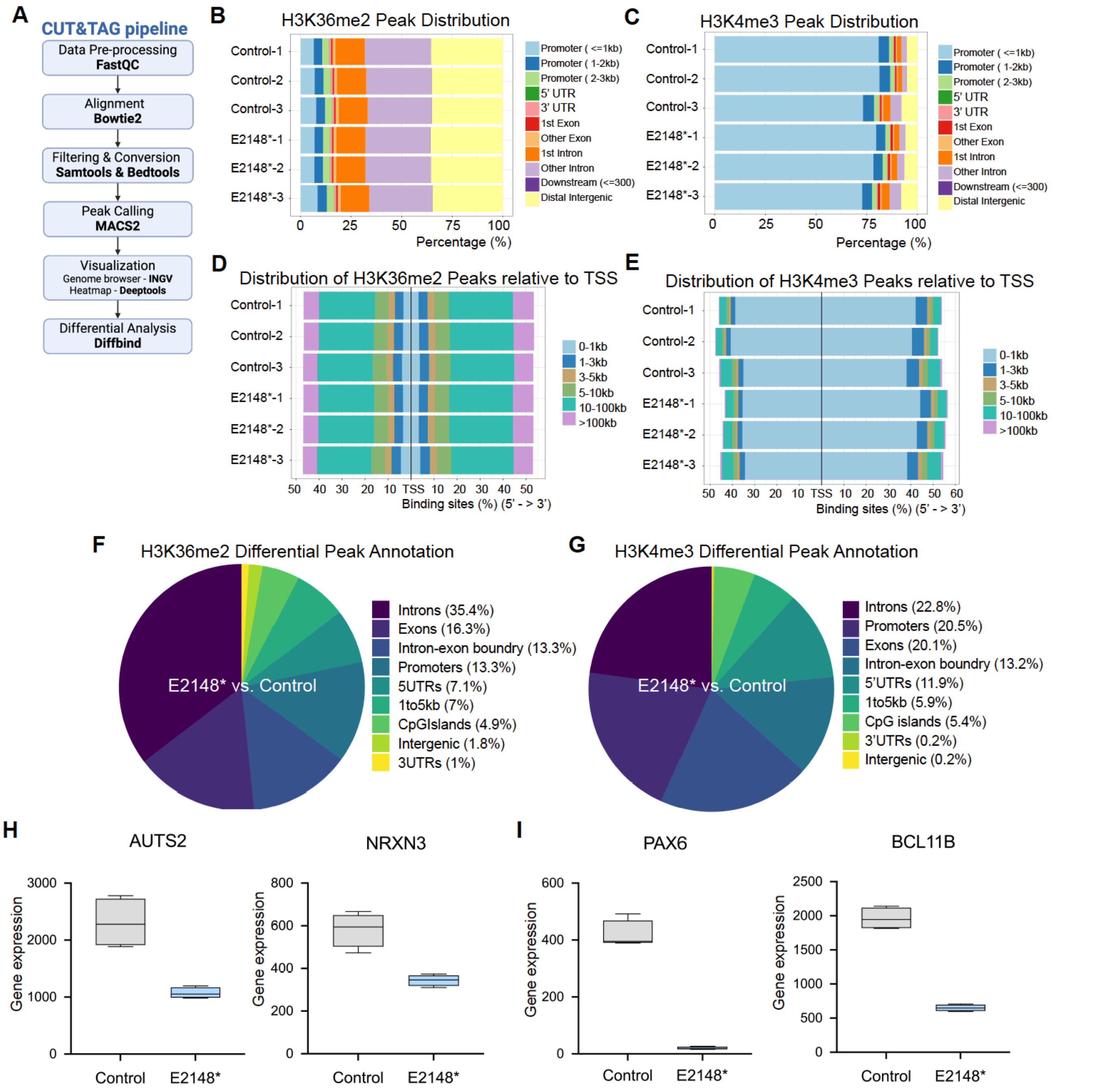


**Supplementary Figure S5: CUT&Tag analysis of E2148* mutant neurons.** (**A**) Bioinformatics pipeline for CUT&Tag studies. (**B-C**) Distribution of all peaks across different genomic regions for H3K36me2 (**B**) and for H3K4me3 (**C**) in control and E2148* neurons. (**D-E**) Distribution of all peaks relative to the transcription start site (TSS) for H3K36me2 (**D**) and for H3K4me3 (**E**) in control and E2148* neurons. (**F-G**) Distribution of differential peaks across different genomic regions is shown for H3K36me2 (**F**) and H3K4me3 (**G**) by comparing E2148* versus (vs.) control neurons. (**H-I**) Box plots showing expression analysis by RNAseq of significant DEGs with differential peaks in H3K36me2 (**H**) or H3K4me3 (**I**) for control (grey) and E2148* (blue) mutant neurons.

**Supplementary Figure S6 relevant to main figure 5.**

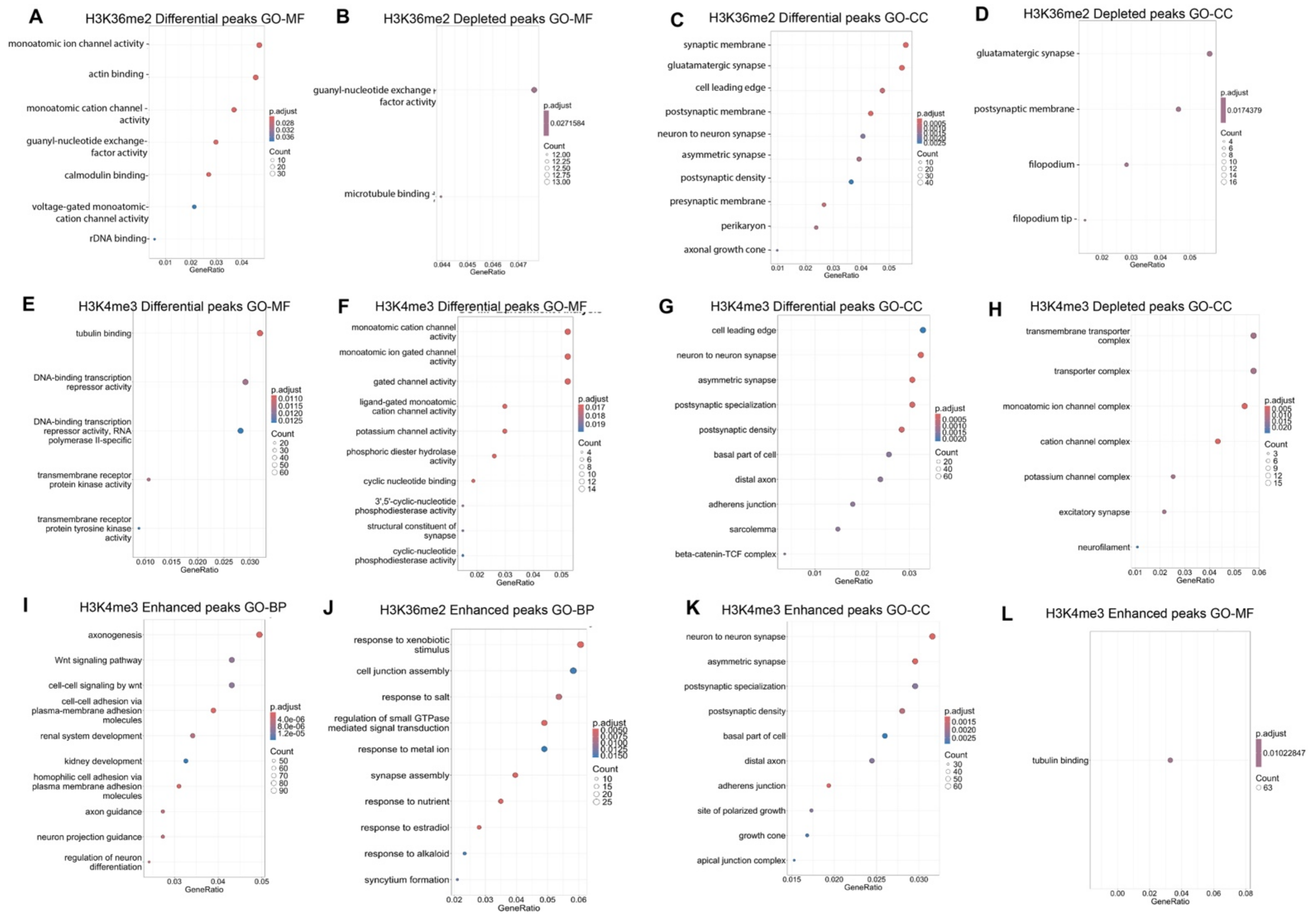


**Supplementary Figure S6. Functional enrichment of H3K4me3 and H3K36me2 differential peaks and mRNA expression levels of representative genes.** (**A-B**) Dot plots showing gene ontology (GO) molecular function (MF) enrichment for H3K36me2 differential peaks (**A**) and depleted peaks (**B**) in E2148* mutant neurons. (**C-D**) Dot plots showing gene ontology cellular compartment (CC) enrichment analysis of genes with differential peaks (**C**) and depleted peaks (**D**) for H3K36me2 in E2148* mutant neurons. (**E-F**) Dot plots showing gene ontology (GO) molecular function enrichment for H3K4me3 differential peaks (**E**) and depleted peaks (**F**). (**G-H**) Dot plots showing gene ontology cellular compartment (CC) enrichment analysis of genes with differential peaks (**G**) and depleted peaks (**H**) for H3K4me3 in E2148* mutant neurons. (**I-J**) Dot plots showing gene ontology biological process (BP) enrichment analysis genes with enhanced peaks for H3K4me3 (**I**) and H3K36me2 (**J**) in E2148* mutant neurons. (**K-L**) Dot plots showing gene ontology biological process (CC) (**K**) and molecular function (MF) (**L**) enrichment analysis for genes with enhanced peaks for H3K4me3 in E2148* mutant neurons.

**Supplementary Figure S7 relevant to main figure 6.**

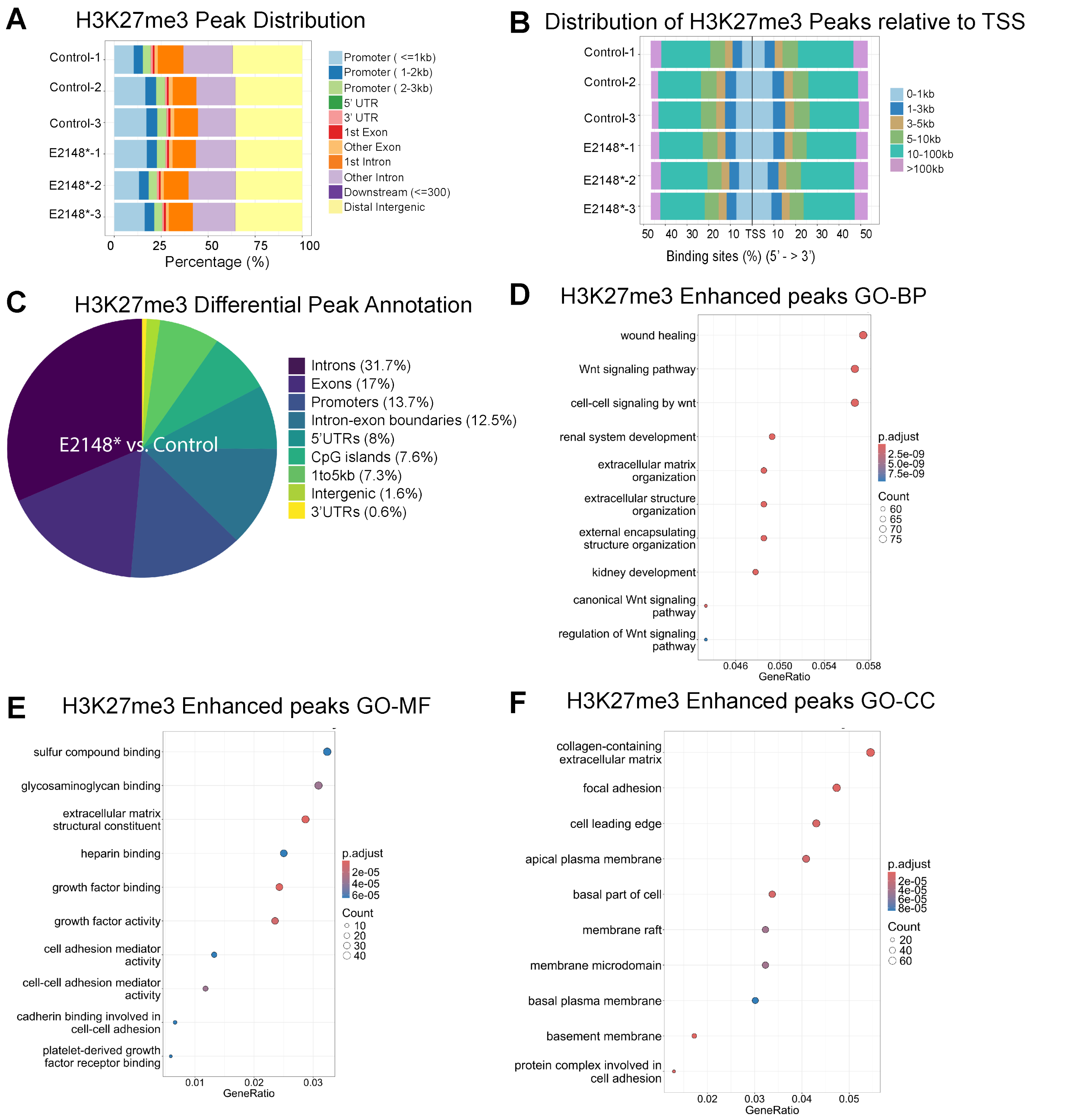


**Supplementary Figure S7. CUT&Tag analysis of H3K27me3 differential peaks.** (**A-B**) Distribution of all H3K327me3 peaks across different genomic regions (**A**) and relative distance to the transcription start site (TSS) for H3K27me2 peaks in control and E2148* mutant neurons. (**C**) Distribution of differential peaks across different genomic regions is shown for H3K27me3 by comparing E2148* versus (vs.) control neurons. (**D-F**) Dot plots showing Gene ontology (GO) enrichment results for genes with enhanced H3K27me3 peaks across biological process (BP) (**D**) molecular function (MF) (**E**) and cellular compartment (CC) (**F**) categories in E2148* mutant neurons.

**Supplementary Figure S8 relevant to main figure 6.**

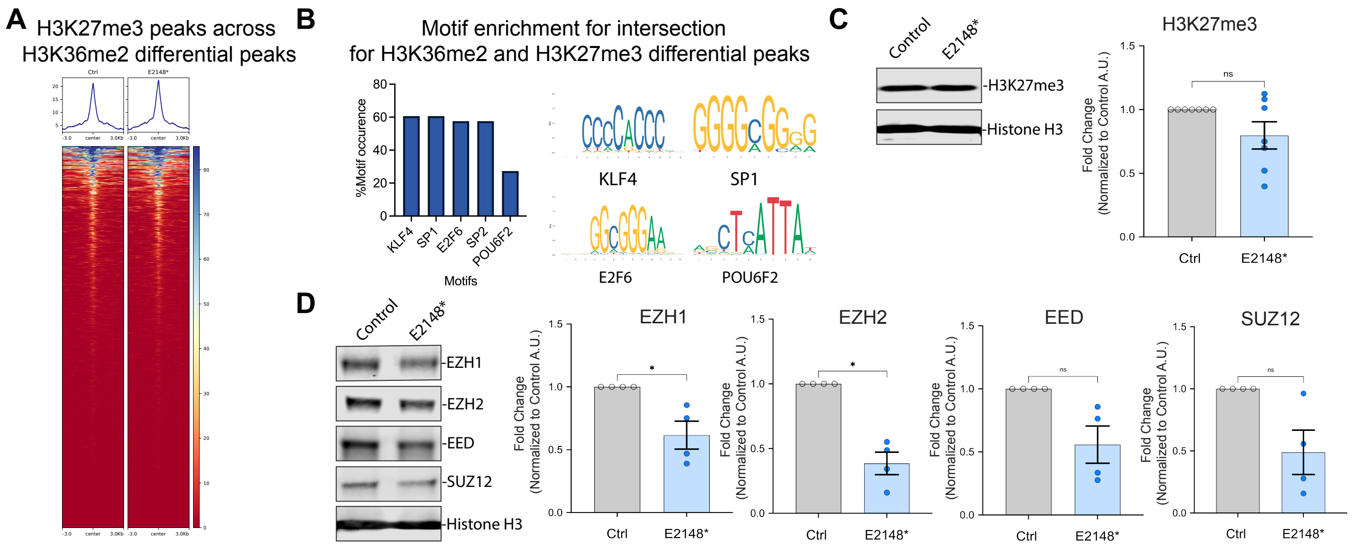


**Supplementary Figure S8. Analysis of H3K27me3 vs. H3K36me2 differential peaks overlap and gene expression of PRC2 subunits.** (**A**) Heatmap showing the distribution of H3K27me3 peaks across H3K36me2 differential peaks shown in a 6Kb interval for control and E2148* neurons. (**B**) Analysis of motif enrichment shown by percent occurrence across regions common for both H3K36me2 and H3K27me3 differential peaks in E2148* mutant neurons. Consensus sequences for the motifs with high percentage occurrence. (**C**) Representative western blot images and quantification for H3K27me3 across control (grey) and E2148* (blue) mutant neurons. (**D**) Representative western blot images and quantification for protein levels of different PRC2 subunits in control (grey) and E2148* (blue) mutant neurons. (**C-D**) Biological replicates are indicated by individual circles. Error bar indicates standard error of the mean. Statistical analysis conducted using unpaired t-tests. *P < 0.05. NS indicates no significant statistical difference.

**Supplementary Figure S9 relevant to main figure 7.**

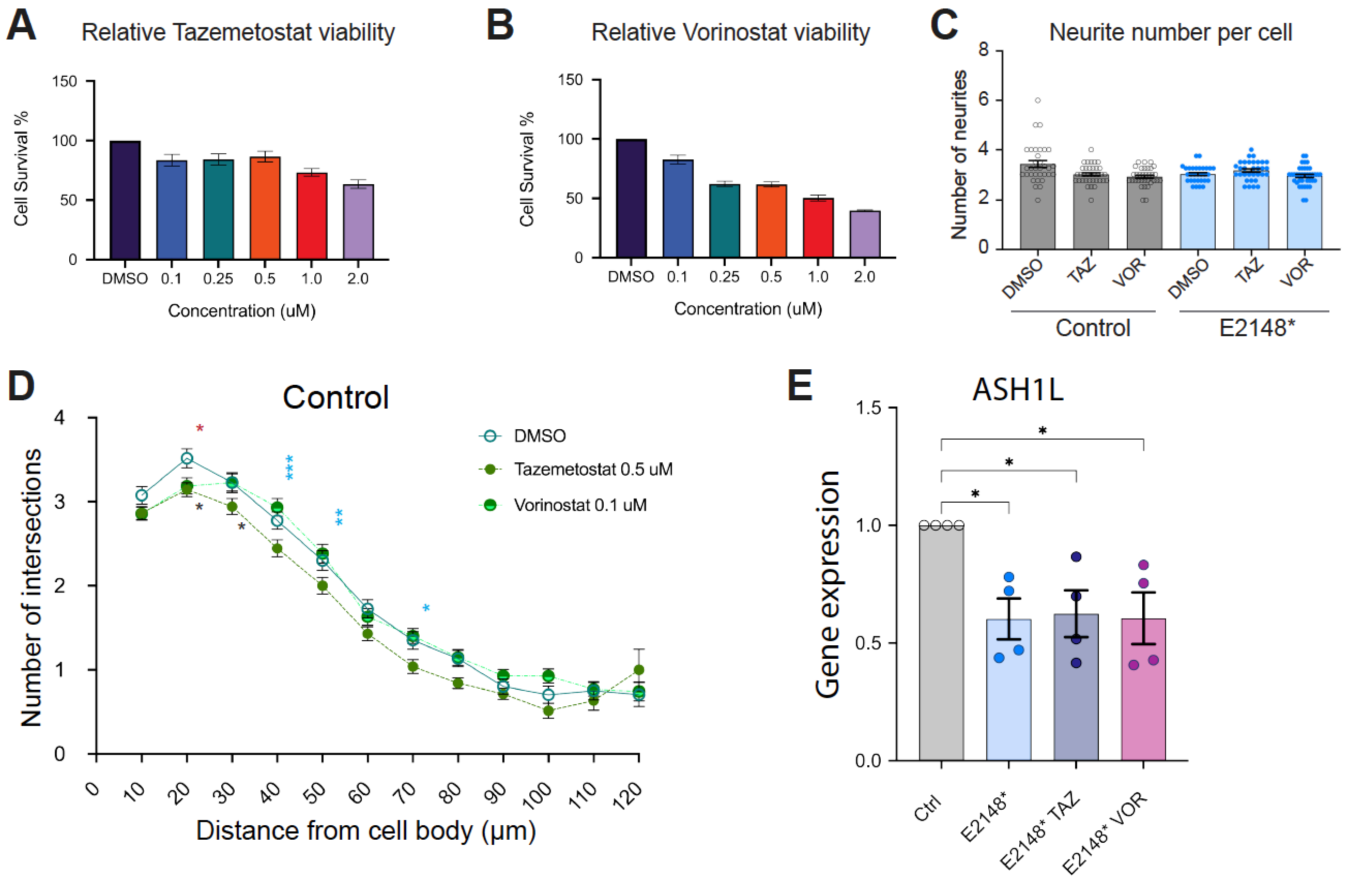


**Supplementary Figure S9. Tazemetostat and Vorinostat effects on cell survival, neurite number and gene expression on control and ASH1L mutant neurons.** (**A-B**) Tazemetostat and Vorinostat effect on cell survival at different concentrations of control neurons. (**C**) Neurite number per cell analysis is shown for at least 3 independent experiments in which we measured at least 30 neurons per experiment for control (grey bar with open circles) and E2148* (light blue bars with solid dark blue circles) neurons treated with either DMSO, Tazemetostat (TAZ) or Vorinostat (VOR). (**D**) Sholl analysis comparing DMSO (open green circle) treatment to TAZ (solid green circle) and VOR (half dark, half light green circle) for control neurons. Statistical analysis was done using a mixed model effect. P values are shown in black (DMSO vs. TAZ), red (DMSO vs. VOR), and blue (TAZ vs. VOR). * P<0.05; **P<0.001; ***P<0.0001. (**E**) Gene expression analysis of ASH1L in response to either TAZ and VOR in control and ASH1L mutant neurons. Error bars indicate standard error of the mean. Statistical analysis was done using one-way ANOVA with correction done using Dunnett's multiple comparison test.

### LEGENDS FOR SUPPLEMENTARY TABLES

**SUPPLEMENTARY TABLE S1**
Table shows sgRNA used to target the ASH1L mutation by genome editing. In addition, amplification primers for genomic and cDNA region amplification as well as sanger sequencing primers are shown.

**SUPPLEMENTARY TABLE S2**
Table shows the TaqMan assay catalog numbers used to conduct qPCR experiments

**SUPPLEMENTARY TABLE S3**
Table shows the DEGs for catalytic mutant vs. control and gene length for DEGs

**SUPPLEMENTARY TABLE S4**
Table shows gene enrichment analysis of DEGs by EnrichR and across GO categories.

**SUPPLEMENTARY TABLE S5**
Table shows ChIP-X Enrichment Analysis of transcription factor binding sites for upregulated and downregulated genes

**SUPPLEMENTARY TABLE S6**
Table shows significant isoform switching events.

**SUPPLEMENTARY TABLE S7**
Table shows DEGs that overlap with SFARI ASD risk genes and gene enrichment analysis across disease pathways

**SUPPLEMENTARY TABLE S8**
Table shows H3K36me2 differential peaks for E2148* mutant neurons vs. controls with associated genomic regions and gene region associations.

**SUPPLEMENTARY TABLE S9**
Table shows H3K4me3 differential peaks for E2148* mutant neurons vs. controls with associated genomic regions and gene region associations.

**SUPPLEMENTARY TABLE S10**
Table shows gene enrichment analysis for H3K4me3 and H3K36me2 differential, depleted and enhanced peaks in E2148* mutant neurons.

**SUPPLEMENTARY TABLE S11**
Table shows motif analysis for H3K4me3 and H3K36me2 differential peaks for E2148* mutant neurons.

**SUPPLEMENTARY TABLE S12**
Table shows H3K27me3 differential peak analysis for E2148* mutant neurons vs. controls with associated genomic regions and gene region associations.

**SUPPLEMENTARY TABLE S13**
Table shows gene enrichment analysis for H3K27me3 differential and enhanced peaks

**SUPPLEMENTARY TABLE S14**
Table shows motif analysis for H3K27me3 differential peaks for E2148* mutant neurons.

**SUPPLEMENTARY TABLE S15**
Table shows motif analysis for regions overlapping in H3K27me3 and H3K36me2 differential peaks for E2148* mutant neurons.
